# Network analysis with either Illumina or MinION reveals that detecting vertebrate species requires metabarcoding of iDNA from a diverse fly community

**DOI:** 10.1101/2022.08.18.504443

**Authors:** Amrita Srivathsan, Rebecca Loh Ker, Elliott James Ong, Leshon Lee, Yuchen Ang, Sujatha Narayanan Kutty, Rudolf Meier

## Abstract

Metabarcoding of vertebrate DNA obtained from invertebrates (iDNA) has been used to survey vertebrate communities, but we here show that it can also be used to study species interactions between invertebrates and vertebrates in a spatial context. We sampled the dung and carrion fly community of a swamp forest remnant along a disturbance gradient (10 sites: 80-310m from a road). Approximately, 60% of the baited 407 flies yield 294 vertebrate identifications based on two COI fragments and 16S sequenced with Illumina and/or MinION. A bipartite network analysis finds no specialization in the interaction between flies and vertebrate species, but a spatial analysis revealed that surprisingly 18 of the 20 vertebrate species can be detected within 150m of the road. We show that the fly community sourced for iDNA was unexpectedly rich (24 species, 3 families) and carried DNA for mammals, birds, and reptiles. They included common and rare ground-dwelling (e.g., wild boar, Sunda pangolin), and arboreal species (e.g., long-tailed macaque, Raffles’ banded langur) as well as small bodied vertebrates (skinks, rats). All of our results were obtained with a new, greatly simplified iDNA protocol that eliminates DNA extraction by obtaining template directly through dissolving feces and regurgitates from individual flies with water. Lastly, we show that MinION- and Illumina-based metabarcoding yield similar results. Overall, flies from several families (calliphorids, muscids and sarcophagids) should be used in iDNA surveys because we show that uncommon fly species carry the signal for several vertebrate species that are otherwise difficult to detect with iDNA.

## INTRODUCTION

Over the last 45 years, forest vertebrate populations have declined by >50% (Green et al., 2020), and numerous species are on the brink of extinction. Many of these are hard to monitor, because they are small, nocturnal and/or arboreal (see discussion in Srivathsan et al., 2019). The only good news is that monitoring can now be facilitated by many new tools. They are complementing traditional field techniques such as transects (e.g., Ang, Ismail, & Meier, 2010), detection of footprints, burrows and feces (Harmsen, Foster, Silver, Ostro, & Doncaster, 2010; Harrington, Harrington, Hughes, Stirling, & Macdonald, 2010), live-trapping (Flowerdew, Shore, Poulton, & Sparks, 2004), and the all-important camera trapping (Welbourne, Claridge, Paull, & Ford, 2020), which yields data that can be analysed semi-automatically using species detection algorithms.

A particularly important recent addition to the tool kit for biomonitoring is the analysis of iDNA: i.e., vertebrate DNA sourced from invertebrates (Calvignac-Spencer et al., 2013; Schnell et al., 2015, 2012). Initially, the main source for iDNA was hematophagous invertebrates such as leeches (Abrams et al., 2019; Schnell et al., 2018; Siddall et al., 2019; Weiskopf et al., 2018), mosquitoes (Hernández-Triana et al., 2017; Kent, Norris, & Feinstone, 2005; Reeves, Gillett-Kaufman, Kawahara, & Kaufman, 2018), phlebotomine sand flies (Kocher et al., 2017), ceratopogonid biting midges (Lassen, Nielsen, Skovgård, & Kristensen, 2011), frog-biting midges (Cutajar & Rowley, 2020), and ticks (Gariepy, Lindsay, Ogden, & Gregory, 2012; Weiskopf et al., 2018). However, another abundant source was found in the form of carrion and dung-feeding insects such as blow- and fleshflies (Calvignac-Spencer et al., 2013; Lee, Gan, Clements, & Wilson, 2016; Owings et al., 2019; Owings, Gilhooly, & Picard, 2021) and dung beetles (Drinkwater et al., 2021). The availability of a wide range of iDNA sources has been an important advantage of the technique because collecting relevant samples often requires little effort and can be integrated into existing field protocols. Some iDNA can be picked up serendipitously during routine surveys (e.g., from leeches), while carrion flies can be conveniently collected with baited traps. Data obtained from iDNA have been very promising. For instance, leech iDNA provided occupancy data consistent with results based on camera trapping (Abrams et al., 2019) and several fly-derived iDNA studies show that the technique can detect more smaller-bodied and arboreal species than camera trapping (Gogarten et al., 2020). Furthermore, museum specimens can yield iDNA (leeches: Siddall et al., 2019) and thus data critical for reconstructing original species ranges and estimating when invasive species arrived.

However, iDNA is currently used mostly as a detection tool for mammals. For example, it is common that only the mammal records are discussed, although iDNA is particularly useful for detecting lizards, snakes, birds, and even fish that are overlooked by many other survey techniques (Lee et al., 2016; Rodgers et al., 2017). In addition, little or no attention is paid to the taxonomic identity of the iDNA carriers (e.g., “carrion flies”), although, for example, standard carrion bait traps attract a large number of species belonging to several Diptera families. The biology of these species is likely to differ which may affect how the newly obtained vertebrate data should be interpreted. An obvious factor is the mobility of the iDNA-carrying fly species, which will impact the spatio-temporal resolution of the iDNA data. For example, some fly species are excellent fliers (Bomphrey, Walker, & Taylor, 2009) which is good news for detecting vertebrates in inaccessible or particularly sensitive sites that can be replete with rare species (e.g., Chambers et al., 2007). Good fliers, however, would be a poor source of information if the vertebrate fauna of two neighboring habitats have to be compared.

We here argue that iDNA should not only be used as a mammal detection technique, but as a tool for understanding species interactions between vertebrate and invertebrate communities. This is feasible because sequencing cost is now so low that large-scale iDNA analyses based on individual invertebrate specimens are feasible (Bush et al., 2020; West et al., 2021) and large amounts of species interaction information can be obtained rapidly (Lee et al. 2022). Indeed, studying species interactions based on iDNA is a logical next step because it not only contributes to our understanding of community ecology, but also helps with sharpening the iDNA tool for vertebrate detection if certain invertebrate species are specialists for certain vertebrate species. The most straightforward improvement over current procedures is the simultaneous species-level analysis of the insect communities that yield the iDNA signal and the vertebrates that are detected. This should be carried out along spatial transects, so that additional information on habitat specialization can be collected. Such an expanded use of iDNA would start with determining the abundance and spatial distribution of fly communities and their iDNA signal. The next step could then be the use of metabolite screens for distinguishing whether the iDNA comes from feces or meat (see Owings et al., 2019, 2021).

In our study, we thus treat the vertebrate community of our field site (a Swamp forest remnant in Singapore) and the community of fly species attracted to our bait (rotten fish) as two sets of species whose interactions are analyzed. In particular, we test whether there is evidence for certain fly species to feed predominantly on certain vertebrate species. This is well known for hematophagous invertebrates (Hernández-Triana et al., 2017; Reeves et al., 2018), but much less clear for carrion flies, although congeneric species can have very different feeding preferences: for instance, the females of *Chrysomya bezziana* attack mostly domesticated animals where oviposition takes place on wounds (Ferrar, 1987). The adult females tend to feed on blood and sera while males only require flower nectar (Norris & Murray, 1964). On the other hand, the congener *C. megacephala* is largely necrophagous and found on carcasses of many vertebrate species (Ferrar, 1987; Norris & Murray, 1964). In addition, vertebrate detection will also be dependent on habitat preferences of carrion fly species. For instance, *Mesembrinella bellardiana* is mostly found in well-protected habitats and would be a poor choice for detecting vertebrates in urbanized environments (Cabrini, Grella, Andrade, & Thyssen, 2013).

However, we here not only argue that iDNA should be used for community ecology, but we also provide the new methods that enable this expanded use by introducing new scalable methods for the study of iDNA from individual insects. Traditionally, iDNA is extracted from blood meals, gut contents, or whole individuals. After DNA extraction, short fragments of identification genes (usually 12S, 16S, cytochrome c oxidase subunit I (COI)) are amplified and sequenced. Sanger sequencing or 454 sequencing (Calvignac-Spencer et al., 2013; Lee, Sing, & Wilson, 2015) were used in the past, but have nowadays been mostly replaced with Illumina sequencing (Drinkwater et al., 2021; Massey et al., 2021).

Arguably, the main limitation of iDNA-based vertebrate monitoring with carrion flies has been the high cost of sample processing. Routine surveys often yield thousands of flies. This means that researchers have to either subsample or pool flies (Gogarten et al., 2020). The latter has a mixed track record, because rare vertebrate signals can be lost by pooling (Calvignac-Spencer et al., 2013; Rodgers et al., 2017). In addition, many pools involve a mixture of fly species, so that this approach cannot yield insights into species interactions. Lastly, many studies extract DNA from fly guts, which first have to dissected laboriously from abdomens (Drinkwater et al., 2021). Depending on the method of DNA extraction, we estimate the consumable cost per specimen to be 1-10 USD given that most studies rely on commercially available DNA extraction kits such as GeneMATRIX Stool DNA purification kit and QIAGEN DNeasy Blood and Tissue kit (Axtner et al., 2019; Calvignac-Spencer et al., 2013; Gogarten et al., 2020; Siddall et al., 2019). These techniques are too expensive for many countries that host most of the biodiversity but lack sufficient funding or resources to process thousands of samples for routine sequencing. More cost-effective methods have only been proposed for blood meals where Whatman four-sample Flinders Technology Associates (FTA) cards (Sigma-Aldrich Corp.,St. Louis, MO) have been used for sample preservation (Reeves et al. (2018) before extracting the DNA using hotSHOT (Truett et al., 2000).

To popularize the study of iDNA, we here introduce a greatly simplified protocol. It skips DNA extraction by generating template through dissolving fly feces and/or regurgitates in water. These digestive fluids can be readily obtained from collecting vials used to house individual flies. We furthermore reduce the amount of time needed to obtain data by exploring the use of nanopore sequencing. The use of ONT sequencing devices would be useful because reliable Illumina sequencing facility are lacking in many biodiverse countries, so that samples are usually exported for sequencing elsewhere (Lynggaard et al., 2019). In addition, the initial set up cost for a MinION based lab is low (<10,000 USD: Srivathsan, Hartop, et al., 2019) and the data can be generated real time which makes MinION sequencers attractive for biodiversity surveys (Blanco et al., 2020; Menegon et al., 2017; Pomerantz et al., 2018). However, a matter of concern has been the high error rates of MinION reads (85-95%: Silvestre-Ryan & Holmes, 2021; Wick, Judd, & Holt, 2019).

Recently, this challenge has been overcome for barcoding individual specimens (Pomerantz et al., 2022; Sahlin, Lim, & Prost, 2021; Srivathsan et al., 2018, 2021), but the analysis of metabarcoding amplicons remained challenging (but see Baloğlu et al., 2021; Davidov et al., 2020; Loit et al., 2019). Currently, it still appears unlikely that complex communities can be characterized reliably using MinION, especially if reference DNA barcodes are not available and the sampled community includes many closely related species. However, MinION sequencing is already suitable for the analysis of metabarcoding data when the species diversity per sample is small, the target species are well represented in publicly available databases, and there is a checklist of expected species for external verification (see recent analysis of seafood products in Ho, Puniamoorthy, Srivathsan, & Meier, 2020).

We here apply a community ecology approach to iDNA to one of the areas in Singapore that most require it; i.e., the largest swamp forest remnant. We analyze the impact of human disturbance on fly-detected vertebrate communities by studying samples collected along a disturbance gradient consisting of ten sites ranging from proximal to a road (80 m) to deeper into the forest (312 m). The study site is heavily impacted by human activity which is a major driver for changes in forest communities (Bryan et al., 2020; Bennett, 2017; Nafus et al.,2013; Clarke et al., 2013). Alterations to vertebrate communities are therefore expected near areas with high human pressure, but studies on the effect of roads have been so far species-specific and used traditional surveillance methods (Bennett, 2017). Use of iDNA provides an opportunity to study the effect of proximity to human disturbance on entire vertebrate communities with the caveat that it still remains unclear how much spatial resolution is lost due to invertebrate movements in the habitat.

Overall, this study thus tests the impact of human disturbance on fly-vertebrate communities. We furthermore analyze the species interaction network between the two communities to understand whether there is any preference or specialization and whether the analysis of many fly species yields more vertebrate records. Thirdly, we demonstrate the effectiveness of a simplified protocol for obtaining iDNA and analyzing it rapidly with Nanopore or Illumina sequencers.

## METHODS

### Sampling

Flies were baited using fish meat of a marine fish (*Rastrelliger kanaguta*) that was obtained from supermarkets in Singapore and allowed to rot for two to three days in an airtight container. In the field, the bait was used to attract flies using a plastic container covered with mesh to prevent flies from landing on the bait. The bait was then placed within a foldable butterfly cage with the zipper left partially open to admit flies. Once flies landed on the inside of the mesh cage, they were caught using 8 ml autoclaved Sarstedt vials. To avoid contamination, new Sarstedt vials were used for each fly and gloves were worn during the collection of flies. Fly trapping was conducted for two hours at each station and between sites the butterfly cage and container were sterilized using 10% Clorox.

The traps were placed at ten different coordinates in the largest swamp forest remnant in Singapore between May to July 2022 (Supplementary Table 1, Fig 1C). The Nee Soon Swamp Forest is a part of the Central Catchment Nature Reserve in Singapore and contains both secondary rainforest and freshwater swamp forest. The forest has an area of 5 km^2^ and has a rich fauna and flora (Lim et al., 2016; Wong et al., 2013; Yeo et al., 2021). The forest is however disturbed and surrounded by multiple roads and a golf course. The traps in this study were placed close to a road to assess if the distance from human disturbance impacts the fly and vertebrate community. In particular, we tested whether dense sampling in the vicinity of a road (Old Upper Thomson Road) can eliminate the need for sampling deep in the forest.

**Figure 1:**
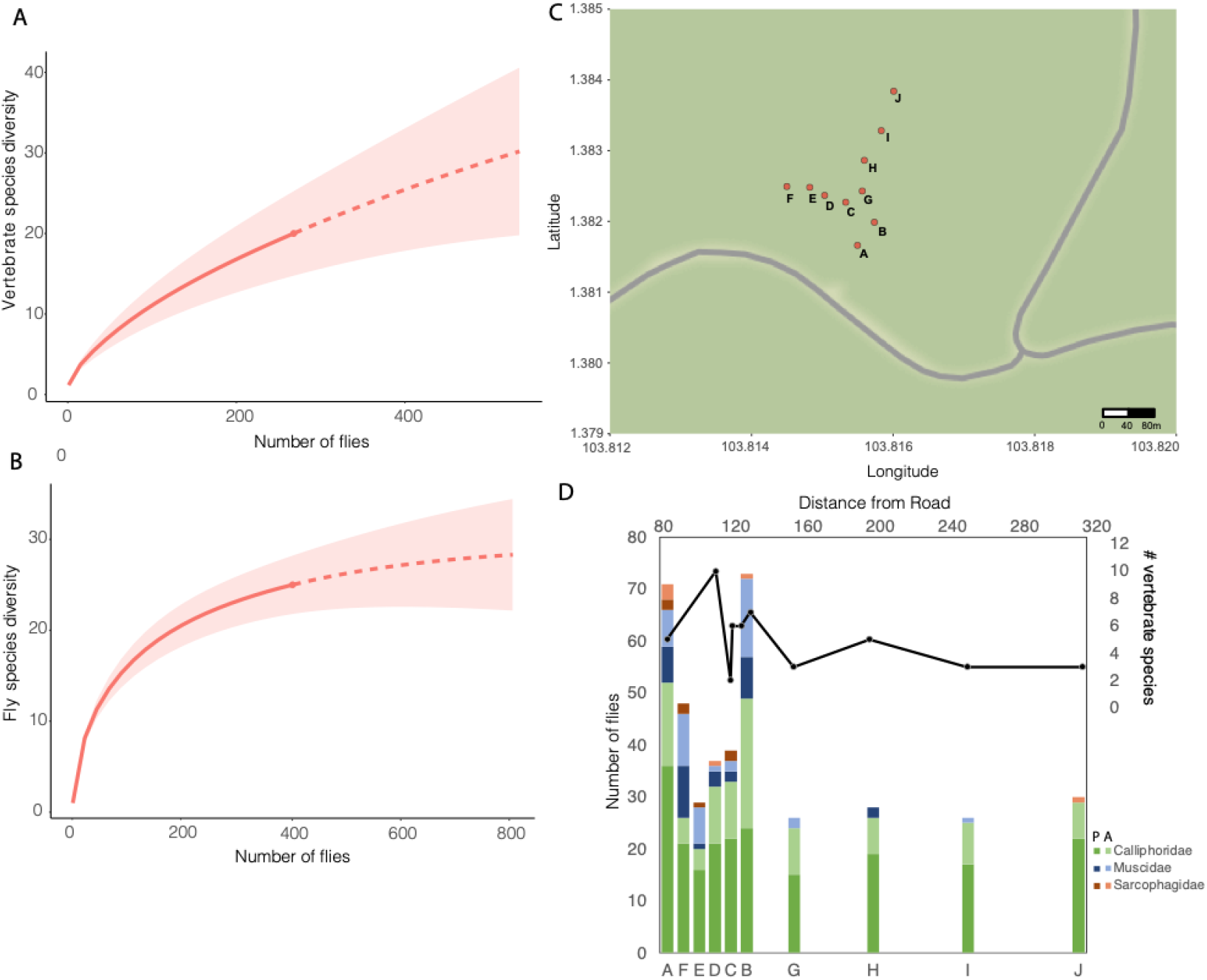
Species accumulation curves for (a) vertebrate and (b) fly diversity. (c) Sampling locations for the study. (d) Vertebrate diversity at each site is shown by the line graph. Bar-graph shows the number of flies with and without vertebrate identifications (P: present, A: absent). The x-axis of the bar graph has been approximately scaled to match the line graph.

### PCR amplifications for fly fecal samples and regurgitates

Flies were transported to laboratory and individual flies were frozen in −30°C after half an hour of arrival of the samples to allow for flies to defaecate or regurgitate food. The estimated transport time was ~30 min and flies were kept alive for at least another 30 min. Overall, the flies were kept alive in vials for at most 2 hours before being frozen. Frozen flies were later removed from the Sarstedt vials by decanting and preserving them in 95% ethanol, while fecal samples and fly regurgitates on the inner walls of the tubes were classified as “brown” (suggesting the presence of fecal samples), “clear” (suggesting the presence of regurgitated digestive fluid), “both”, or “empty” (if the tube contained nothing visible). After noting the appearance of fly excretions, 20–50 μL of molecular grade water was pipetted onto the fly exudes to dissolve them. The amount of water was determined based amount of regurgitates or feces observed. If vials appeared to be empty or contained amount of clear liquid similar to condensation, 20 μL of molecular grade water was used to wash the inner surface of the tubes. For other samples having bigger drops of regurgitates or feces, 50 μL of molecular grade water was used.

PCR amplifications were done using tagged primers to allow for multiplexing large number of samples on Next Generation Sequencing platforms. We used three different primer pairs in this experiment. One targets the 313-bp COI minibarcode of metazoans, m1COlintF: 5’-GGWACWGGWTGAACWGTWTAYCCYCC-3’ (Leray et al., 2013) and modified jgHCO2198: 5’-TANACYTCNGGRTGNCCRAARAAYCA-3’ (Geller, Meyer, Parker, & Hawk, 2013); a second pair targets the 244-bp COI minibarcode of vertebrates, Mod_RepCOI_F: 5’-TNTTYTCMACYAACCACAAAGA-3’ and VertCOI_7216_R: 5’-CARAAGCTYATGTTRTTYATDCG-3’ (Reeves et al., 2018); the last pair targets the 93-bp 16S minibarcode of mammals,16Smam1: 5’-CGGTTGGGGTGACCTCGGA-3’ and 16Smam2: 5’-GCTGTTATCCCTAGGGTAACT-3’ (Taylor, 1996). The primers were tagged with 13-bp indices that are suitable for both nanopore and Illumina sequencing (Srivathsan, Hartop, et al., 2019). PCRs were conducted using 10X dilutions of the dissolved material, applying the following PCR recipe: 11 μL of mastermix from CWBio, 2 μL of 1 mg/mL BSA, 2 μL of 10 μM of each primer and 5 μL of DNA. The following PCR conditions were used for the 313-bp COI minibarcode: 3 min initial denaturation at 94 °C followed by 35 cycles of denaturation at 94 °C (0.5 min), annealing at 45 °C (1.5 min), extension at 72 °C (0.5 min), and followed by final extension of 72 °C (3 min). The PCR conditions for the 244-bp COI minibarcode were as follows: 3 min initial denaturation at 94 °C followed by 40 cycles of denaturation at 94 °C (40 s), annealing at 48.5 °C (2 min), extension at 72 °C (0.5 min), and followed by final extension of 72 °C (3 min). PCR conditions for the 93-bp 16S minibarcode were as follows: 10 min initial denaturation at 94 °C followed by 40 cycles of denaturation at 94 °C (12 s), annealing at 59°C (30 s), extension at 70 °C (25 s), and followed by final extension of 72 °C (7 min). Three independent PCR replicates were performed for each sample, each of the three replicates were run in separate PCR plates with each plate containing a negative.

Nine independent amplification-free DNA libraries were prepared for Illumina sequencing. Each library consisted of samples from a single PCR replicate and three libraries each were prepared per primer pair. Libraries were prepared using VAHTS Universal Pro DNA Library prep kit for Illumina^®^ ND608. Library preparation and sequencing was outsourced to GeneWiz (From Asenta Life Science) using Illumina NovaSeq to obtain 250X2 bp sequences for the 313-bp COI amplicon, and 150X2 bp sequences for the 244 bp COI amplicon and 16S amplicons.

In order to assess the performance of nanopore sequencing, we also sequenced one PCR replicate for the amplicon pools of all three primer pairs with MinION. The DNA library was prepared using SQK-LSK110 Ligation Sequencing Kit (Oxford Nanopore Sequencing) for the 313-bp COI amplicon pool and the SQK-LSK112 kit for the 244-bp COI and 93-bp 16S amplicon pools, using recommended protocols with the modifications as follows: We excluded the FFPE DNA repair mix in the end-repair reaction and instead the reaction consisted of 50 μl of DNA, 7 μl of Ultra II End-prep reaction buffer (New England Biolabs), 3 μl of Ultra II End Prep enzyme mix (New England Biolabs). An additional modification was the use of 1X ratio of Ampure XP beads accounting for the short fragments lengths sequenced in this study. Sequencing was conducted on three separate flowcells for each amplicon pool (313-bp COI: R10.3 flowcell, MinION Mk1B; 244-bp COI: R10.4 flowcell, MinION Mk1B; 93-bp 16S: R10.4 flowcell, MinION Mk1C). The 313-bp COI MinION run yielded only 4 million sequences, which is lower than usually obtained and we therefore obtained an additional ca. 2 million sequences by reusing the same flowcell and another one after washing. Note that the second flowcell had previously not been used for the targeted 313 bp COI fragment.

### Data analysis

The FASTQ files containing Illumina sequences were processed using the OBITOOLs pipeline (Boyer et al., 2016). Prior to this, for 313-bp COI and 16S datasets, paired ends reads were merged using PEAR (v0.9.8) (Zhang et al. 2014). For 244-bp COI dataset that was sequenced using 150 bp sequencing technology, we merged the reads end to end to with internal padding of Ns using bbmap (Bushnell, 2014). Next, *ngsfilter* was used to demultiplex the sequences allowing for 2 bp mismatch at the primer sequences. Unique sequences were then obtained using *obiuniq*. Short sequences (≤300 bp for 313-bp COI fragment and ≤200 bp for 244-bp COI fragment and ≤50 bp for 16S) and singleton sequences were excluded using *obigrep*. *obiclean* was used to identify erroneous sequences. Non erroneous sequences, termed “head” and “singleton” were further processed. Lastly, only sequences present in at least 2 replicates were retained for further analysis.

These sequences were matched to the NCBI *nt*-database using BLASTN with minimum e-value of 1e-10. BLASTN matches were parsed using *readsidentifier* (Srivathsan, Sha, Vogler, & Meier, 2015). Here, for any sequence, the matches with the best identity are retained and then classified based on the lowest common ancestor algorithm. Here the hit-length parameter was set to >300 bp hit length for 313-bp COI fragments and >50 bp for 16S. >50 bp hit-length used for 244-bp COI fragment which was split back to single end reads prior to BLAST matches. Sequences having best matches to chordates were then extracted. Afterwards, we assessed total number of vertebrate reads for each of the markers (313-bp fragment of COI: 1,695,815, 244-bp fragment of COI: 14,338,243 and 16S: 183,337,425) and found it to vary widely. Using these vertebrate coverages, we established proportional count cut-offs. They were 2 (for 313-bp COI fragment), 20 (for 244-bp COI fragment) and 250 (for 16S). Based on these filters, the negatives were clean with the exception of half a plate of 16S amplicons where the negatives yielded a *Macaca fascicularis* hit. We thus excluded *Macaca* identifications based on 16S fragments for this half plate.

Due to the use of different genes and sequencing platforms, we explored multiple identity cut-offs for obtaining species identifications. For all Illumina data, we applied the following filters: Firstly, we excluded human reads. These comprised of 23% of 313-bp COI, 66.7% of 244-bp COI fragment and 6.88% of 16S fragment. Human reads could be contaminations or real signals, but are not of interest for vertebrate surveillance. Removal, however, could affect our community-level analysis because one species (*Homo sapiens*) that occasionally is likely to leave iDNA signal in the forest has been removed. Secondly, when congeneric matches were present in the same fly, the sequence with higher identity was accepted as the accurate species ID and ties were broken based on counts. This was mostly observed in the where the genus *Macaca* had very high read counts. This approach was applied because the likelihood of a single fly visiting congeneric species of vertebrates in Singapore is very low. In addition, most congeneric matches included one Singaporean and one non-Singaporean species (*Macaca mulatta* with *Macaca fascicularis*). For COI, we furthermore conducted a translation-check to ensure the sequences used for identifications were not paralogs. Lastly for 16S, due to the high sequencing depth, it was often observed that the same fly sample yielded identifications to members of the same mammal family (e.g. genus *Macaca* with *Mandrillus*). Similar to criterion for genus in COI, we accepted the identification with the highest identity match, and in case of ties, broke the tie with the highest count. With regard to identity levels, we used >=95% for genus and >97% for species level matches with COI. For 16S, we ignored any identification below 97%. In a few cases with genus-level matches, we made additional tentative species identifications based on checklists whenever Singapore only has one species in the genus (see legend for Table 1).

**Table 1.** Vertebrates identified using metabarcoding of fly fecal samples and regurgitates.

| Species | Common Name | Number of flies | Endangered Status / Abundance<sup>3</sup> |
| --- | --- | --- | --- |
| **Mammals** |  |  |  |
| *Callosciurus notatus* | Plantain Squirrel | 1 | LC |
| *Canis lupus* | Dog | 2 | Pet? (introduced via pet trade) |
| *Felis catus*<sup>1</sup> | Cat | 1 | Pet? (introduced via pet trade) |
| *Galeopterus variegatus* | Colugo | 1 | NT |
| *Macaca fascicularis* | Long tailed macaque | 240 | LC |
| *Manis javanica*<sup>1</sup> | Sunda Pangolin | 1 | CR |
| *Maxomys rajah* | Brown Spiny Rat | 1 | VU |
| *Paradoxurus hermaphroditus*<sup>2</sup> | Common palm civet | 7 | LC (*musangus*)/ NA (*hermaphroditus*)* |
| *Presbytis femoralis* | Raffles' banded langur | 3 | CR |
| *Rattus sp.*<sup>1</sup> | Asian house or brown rat | 1 | NA |
| *Rusa unicolor* | Sambar Deer | 5 | NA |
| *Sus scrofa*<sup>4</sup> | Eurasian Wild Boar | 39 | LC |
| *Tupaia glis* | Common treeshrew | 1 | LC |
| **Birds** |  |  |  |
| *Leptocoma brasiliana* | Van Hasselt's Sunbird | 1 | C |
| *Gallus gallus*<sup>4</sup> | Jungle Fowl | 1 | C |
| *Mixornis gularis*<sup>2</sup> | Pin-striped Tit-babbler | 1 | C |
| *Pycnonotus plumosus* | Olive-winged Bulbul | 1 | C |
| **Reptiles** |  |  |  |
| *Eutropis multifasciata* | Common Sun Skink | 1 | LC |
| *Hemidactylus frenatus* | Common House Gecko | 3 | LC |
| *Malayopython reticulatus* | Reticulated Python | 2 | LC |
| **Others** |  |  |  |
| *Dama dama* | European fallow deer | 1 | Food? |
| *Macropus sp.* | Kangaroo | 1 | Zoo? |
| *Rasbora borapetensis* | Red Tailed Rasbora | 1 | Unknown |
| *Bos taurus* | Cow | 1 | Food? |
| *Ovis sp.* | Sheep | 3 | Food? |

1= Ambiguous identification to *silvestris/catus*, for *Felis*, *javanica/pentadactylus* for *Manis*, and *Rattus tanezumi/norvegicus/hoogerwerfi*. For labelling, we have used *catus* throughout for *Felis*. *Manis* is identified to *javanica* based on Singapore checklist. Similarly *Rattus* is identified to either *tanezumi* or *norvegicus*.
2= ID retained as *hermaphroditus* as BLAST match was at 96.7%. However, *musangus* lacks COI. Identity for *Mixornis* was 95.79% to *flavicollis* and 95.47% to *gularis*, and species identification was retained as *gularis* given it is the only known species of *Mixornis* in Singapore.
3= LC= Least concern, NT= Near Threatened, CR: Critically Endangered; A= Abundant, C=Common, NA= Not available. classifications based on Singapore Red Data book List: https://www.nparks.gov.sg/biodiversity/wildlife-in-singapore/species-list.
4= *Gallus gallus* and *Sus scrofa* are retained as free-ranging due to prevalence of junglefowl and wild boar in the region. However domestic sources cannot be ruled out.

Nanopore data were analyzed by first basecalling the reads in Guppy using the super accuracy model. The resulting fastq files were demultiplexed using ONTbarcoder (v 0.1.9) (Srivathsan et al., 2021). The demultiplexed fasta files were matched against NCBI’s *nt* database using BLASTN with e-value parameter set at 1e-10. The BLAST matches were processed using *readsidentifier* to obtain the closest match under the hit-length criterion of 300-bp for 313-bp fragment of COI, 200 bp for 244 bp fragment of COI and 75 for the ~93 bp fragment of 16S and minimum identity match of 90 and 95%. We eliminated congeneric matches (or confamilial matches for 16S) that were present at lower abundances than the main species level signal. For those analyses where the identifications from all three markers were merged, we recorded a genus-level identification if there were conflicts between 2 species level matches (e.g., *Macaca fascicularis* using COI, and *Macaca mulatta* using 16S as *Macaca sp.*). Similarly, if one marker yielded a genus- and another a species-level match, we accepted the genus-level match (*Macaca fascicularis, mulatta* for 16S and *Macaca fascicularis* for COI, as *Macaca sp.*). All identifications based on singleton sequences were discarded. Furthermore, identifications with counts up to the maximum count of vertebrate identifications in negatives were excluded (>7 for 16S 95%, >10 for 16S 90%). To keep the comparison between Illumina and MinION sequencing consistent, neither analysis used the translation check and we only compared the identifications that were obtained for the same replicate.

### Species composition of fly community

We used DNA barcodes to characterize the fly community caught by the traps by applying the MinION based barcoding pipeline described in Srivathsan *et al*. (2021). Fly legs were removed and DNA extracted using 10–15 μL of hotSHOT (Truett et al., 2000). The resulting extract was amplified for the 658-bp COI barcode using HCO2198 and LCO1490 primers (Folmer, Black, Hoeh, Lutz, & Vrijenjoek, 1994) using protocols described in Srivathsan et al. (2021). The primers were tagged with 13-bp sequences suitable for MinION sequencing. The PCR products were pooled in equal volumes and purified using Ampure XP beads (Beckman Coulter). For MinION based barcoding, a library was prepared using SQK-LSK110 Ligation sequencing kit and the products were sequenced in a used MinION R9.4.1 flowcell. Basecalling was conducted using Guppy under the super accuracy model. The resulting sequences were processed using ONTbarcoder (v0.9.1) (Srivathsan et al., 2021) in order to obtain DNA barcodes using 200X coverage as threshold. The resulting barcodes were matched with GBIF’s sequence-id engine (https://www.gbif.org/tools/sequence-id) in order to identify the insects. All identifications with >=97% sequence identity were accepted at species level. Afterwards, the sequences were aligned using MAFFT v7 (Katoh & Standley, 2013) and clustered at 3% using objective clustering in order to determine species diversity and abundance. For four unidentified clusters, genus level identifications were made based on morphology.

### Statistical analysis

The association between fly and vertebrate species was studied in terms of diversity and specialization. In order to understand how fly sampling impacts vertebrate diversity, we treated flies as sampling units and identified vertebrates as the species. Species richness was estimated using *iNEXT* (Chao & Jost, 2012) in R (R Core Team, 2021). We also estimated the fly species richness, where each 3% mOTU was considered a fly species. In order to examine the interaction between fly and vertebrate species, a species interaction network was plotted using *bipartite_D3* in R. Lastly, to assess if fly family membership impacts the diversity of vertebrates detected, Shannon diversity indices were calculated using the package *vegan* for the diversity of vertebrates detected by each site and family (Calliphoridae, Muscidae, Sarcophagidae).

To assess specialization, we followed the approach of Hoenle et al. (2019) and Wehner et al. (2016) described in Blüthgen, Menzel and Blüthgen (2006). Network-level specialization and species level specialization were measured using two-dimensional Shannon entropy 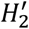 and the Kullback–Leibler distance *d*’, respectively. Both metrics range from 0 to 1 with values closer to 0 suggesting low specificity and values closer to 1 suggesting high specificity. The species interaction network obtained by metabarcoding was compared against null models with fixed marginal totals using Patefield algorithm (“r2dtable”). 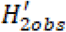 was compared against 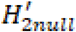 obtained from 10,000 randomizations. The p-value was obtained as the proportion of randomizations yielding 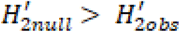. The analysis was conducted on combined network obtained from all three metabarcoding markers and each marker separately, at both species and family levels. All analysis were conducted using R package “bipartite”.

Next, we studied if distance from human disturbance (road) affects vertebrate diversity. The nearest distance from the road was determined using Google Maps by moving the points along the road and assessing the shortest distance. We then established how many vertebrate species are gained as sites further from the road are sampled. To determine whether the observed gain was expected based on the number of flies sampled, we used a newly written script to randomize the placement of specimens in different coordinates (10 coordinates in total, 100 randomizations) while keeping the fly abundances containing vertebrates per site constant. This yielded a randomized gain curve with standard deviations for each sampling site. We furthermore used linear models to test whether the number of vertebrates detected is explained by distance from road and number of flies sampled.

Lastly, we tested whether the probability of detecting a vertebrate species in a fly is influenced by the nature of sample (feces, regurgitate, other trace DNA), sex of the fly, its family membership, or interactions. Tests were conducted separately and one model was implemented which accounted for interactions between the terms. For each test, a binomial logistic regression model (link = “logit”) was implemented using *glm* in R.

## RESULTS

### Vertebrate detections using fly feces and regurgitates

We sampled 435 flies across the ten sites (Supplementary Table 1). Of these, 406 yielded COI barcodes and could be grouped into 25 Operational Taxonomic Units (OTUs) at 3% clustering threshold. Two additional specimens could be identified based on the metabarcoding experiment where the 313-bp COI fragment was amplified. The metabarcoding of the 435 flies using Illumina sequencing yielded 30-40 million sequences per library. Demultiplexing resulted in an average sample coverage of 61,028 sequences per sample per replicate for the 313-bp fragment, an average of 203,496 sequences per sample per replicate for the 244-bp fragment of COI and 147,821 sequences per sample pe replicate for 16S across the 435 samples. For the negatives, we obtained an average of 17,987 sequences for 313-bp COI, 129,073 sequences for 244-bp COI and 7,731 sequences for 16S per replicate. The high read counts for negatives of 244-bp COI fragment is due to co-amplification of bacterial DNA by this primer. After filtering based on read counts, and retaining sequences present in multiple replicates, only one negative corresponding to half a plate of 16S PCRs revealed a non-human vertebrate match (*Macaca*). *Macaca* matches based on 16S were excluded for this plate.

BLAST matches of the metabarcoding data reveal 25 vertebrate species including 18 mammals, 4 birds and 3 reptiles. These were spread across 273 flies, 222 of which yielded a single vertebrate identification. Another 44 gave matches to two different vertebrates while five gave matches to three vertebrates. The most common species was long-tailed macaque (*Macaca fascicularis; n=*240) and wild boar (*Sus scrofa;* n=39), common palm civet (*Paradoxurus hermaphroditus;* n=7) and Sambar deer (*Rusa unicolor;* n=5) while sixteen vertebrates were singleton observations. Notable species on our detection list are four Critically Endangered or Near Threatened species, including Raffles’ Banded Langur (*Presbytis femoralis;* n=3; CR), Sunda Pangolin (*Manis javanica;* n=1; CR), Malayan colugo (*Galeopterus variegatus;* n=1; NT) and the Brown Spiny Rat (*Maxomys rajah;* n=1; VU). The above list of 25 species excludes four *Macaca sp*. and *one Macaca mulatta* match which were retained despite the various filtering criteria. Given that it is unclear if they should be treated as same species as *M. fascicularis*, we excluded these matches. Five of the 25 species are likely to be not from free ranging organisms and are excluded from analysis unless mentioned otherwise: three are known human food (European fallow deer: *Dama dama*, cow: *Bos taurus*, and lamb/sheep: *Ovis sp.*). We also excluded from analysis one species known to exist in the Singapore zoo (kangaroo*: Macropus sp.*). Note that the zoo is 3.6 km from our sampling sites and it cannot be ruled out that carrion flies could travel such a distance (Lee et al., 2015). Lastly we excluded the only aquatic species match (*Rasbora borapetensis*) although it is an introduced species commonly found in Singapore. On the other hand, we retained *Gallus gallus* and *Sus scrofa* in the list of free-ranging animals given the widespread presence of junglefowl and wild boar in the region. Domestic sources for both species cannot be excluded, but it should be noted that there are no pig farms in Singapore and poultry is raised in indoor facilities. Overall, our analyses are thus based on 435 flies (408 with barcodes) yielding 20 validated vertebrate identifications.

### Flies as vertebrate detectors

Vertebrates were identified in 62% of the barcoded flies (254/408) that belonged to 20 species. Individuals of 23 fly species yielded vertebrate identifications. They belonged to three families: Calliphoridae (# total = 316; # with vertebrate identifications = 213), Muscidae (# total = 78; # with vertebrate identifications = 33) and Sarcophagidae (# total = 13; # with vertebrate identifications = 7). One identification was made from a fly species identified as *Leucophenga sp* but the morphological examination of the voucher suggested it is a Micropezidae. This specimen yielded a vertebrate match to *Macaca fascicularis*, but was excluded from further network analysis because morphological and molecular identification did not agree. The vast majority of the flies were female (80.6%). The species accumulation curve for flies suggests that more sampling would further increase the number of species attracted to the bait by *ca*. 30 species (Figure 1). The Chao1 estimate is similar (29.5) but has a large confidence interval (25.8-49.8). Increased sampling of flies will in turn lead to a larger number of vertebrate identifications which is estimated to reach 56 (CI: 27.7-188.1). However, sampling large numbers of flies is not directly correlated with greater vertebrate diversity in all sites. We find that the number of vertebrates species across the sites is variable (2-10 species, Figure 1D: line graph) and this variability does not follow the disturbance gradient (Figure 1D).

The species interaction network between fly hosts and vertebrate species reveals that the calliphorid *Chrysomya megacephala* yields the largest number of interactions (Table 2, Figure 2, video: https://youtu.be/ktNq2slO9pk). This species is associated with 15 different species across mammals, birds and reptiles (13 excluding the “other” vertebrate identifications). Another species of *Chrysomya, C. rufifacies* is associated with seven different mammals (three excluding the “other” identifications). The most common mammal, long-tailed macaque, was associated with 21 different fly species while wild boars are connected to ten fly species. In terms of vertebrate diversity, calliphorid and sarcophagid flies reveal similar diversity, while muscid flies (largely belonging to genus *Atherigona*) detect a lower diversity of vertebrates (Shannon Index: Calliphoridae: 1.04, Muscidae: 0.84, Sarcophagidae:1.21, including non-free ranging animals). It should be noted that the sample sizes for the flies are very different such that total number of vertebrate species detected by Calliphoridae, Muscidae and Sarcophagidae are 20, 4, and 4.

**Figure 2.**
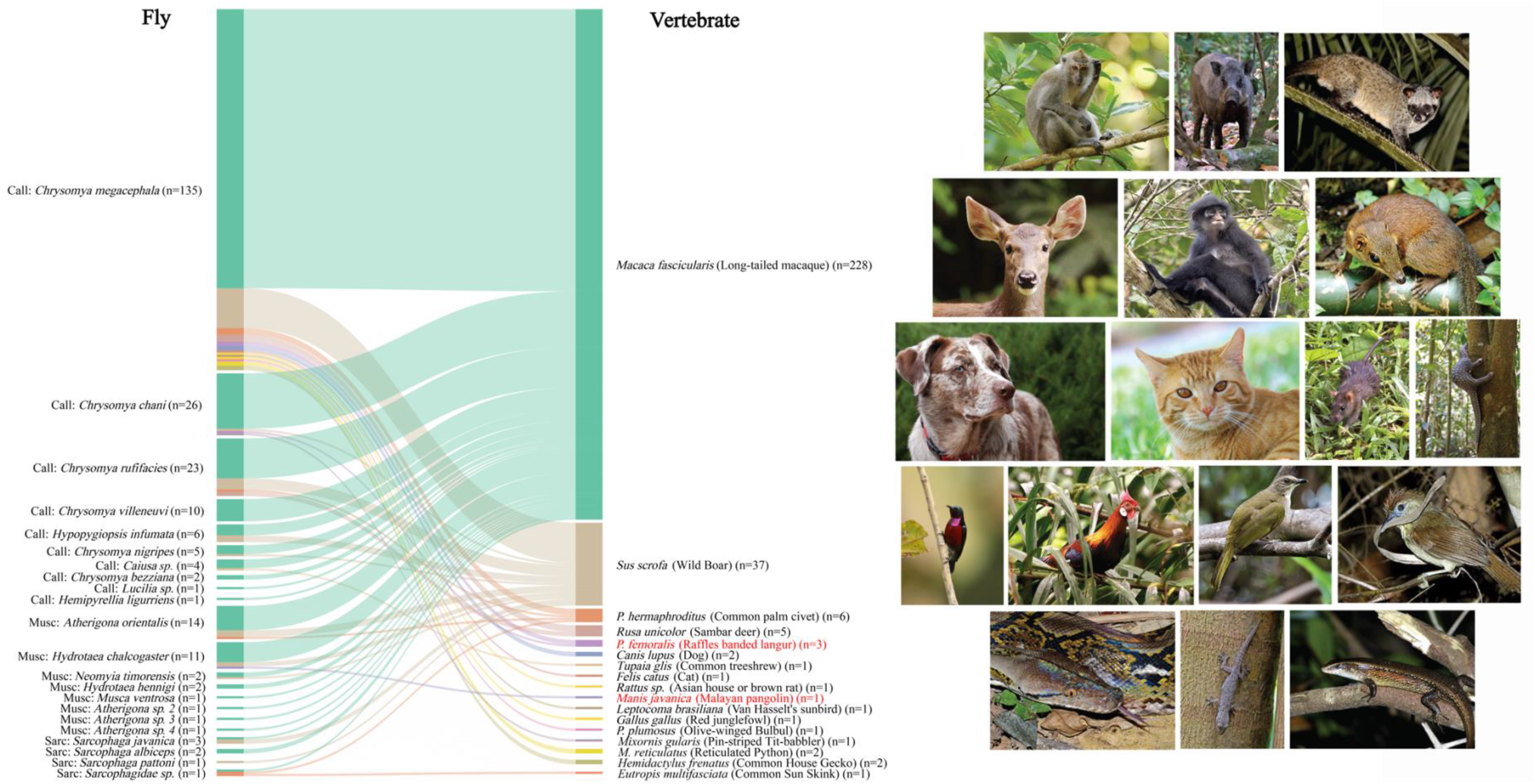
Bipartite graph representing the vertebrate species detected from flies using all the identifications made by all three markers. The values in brackets represent the total number of individuals (flies)/detections (vertebrates) for the species involved in the interactions. Highlighted in red are critically endangered species. All images except dog, cat and budgerigar are taken from Ecology Asia (credit: Nick Baker, permission obtained). Cat: Kala King, dog: Alan Levine.

**Table 2:**
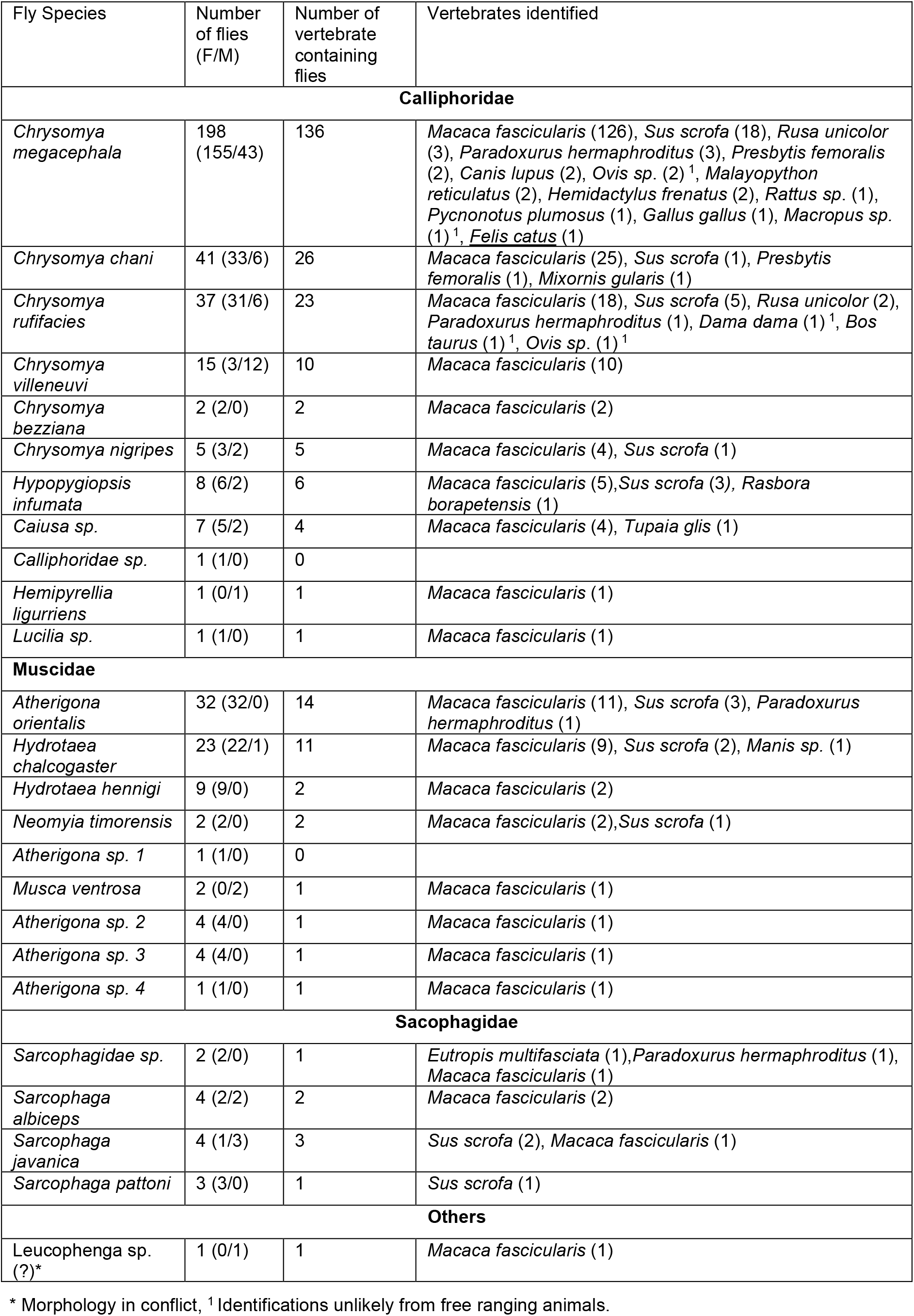
Flies baited using fish baits in and their associations

Network level analysis reveals a generalized network with Shannon diversity for combined network found as close to 0 (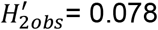 for species level, 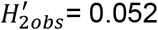 for family level). Comparison with null models places this within random-distribution (P=0.712 for species level and P=0.333 for family level). When Kullback–Leibler divergence *d’* values are examined, 18 species have d’value <0.1 and two had *d’* value between 0.1-0.2 (*Caiusa* sp. and *Sarcophaga javanica*). Two species of *Sarcophaga* had the highest *d’* values (*Sarcophaga sp. d’=* 0.45 and *Sarcophaga pattoni*, d’= 0.34), but both species were rare. The family level network revealed *d’* value <0.1 for calliphorids and muscids and slightly elevated have *d’* for sarcophagids (0.15). When individual metabarcoding markers were examined, we also found generalized networks for 16S and 244-bp fragment of COI. The 313-bp fragment however yielded network with some partitioning such that 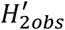 that is higher than most 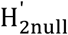 (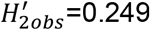, P=0.034).

### Effect of Distance from Road

Sampling was conducted at a range 82-312 m distance from the road to determine whether sampling has to involve sites within the swamp forest, which is often inaccessible due to flooding (Figure 1). We show that within ~150 m from the road, 18 of the 20 vertebrate species are detected (Figure 3). This includes all the threatened or endangered species. The only species missing are reticulated python and a rat species (either Asian house rat *Rattus tanezumi* or brown rat *R. norvegicus*), which was detected deep in the forest, but is known to occur all over Singapore including many urban sites. Randomization of the data across sites suggests that the observed recovery curve is very similar to the predicted curve. This is in line with two linear models that revealed no significant association between the number of vertebrate identifications and distance from the road (vertabundance ~ Distance; p=0.15, R^2^=0.24) and a modified model that included fly abundance as a variable (vertabundance/flyabundance ~ Distance; p=0.59, R^2^=0.039). However, as expected vertebrate abundance is closely correlated with number of flies sampled (p= 2.34e-05, R^2^=0.90), and this model’s fit is very similar to one where the interaction between distance and vertebrate abundance is accounted for (vertabundance~Distance* flyabundance: p=0.0008, R^2^=0.93). Similar patterns are found when assessing diversity of vertebrates: there is no relationship between vertebrate species richness and distance from the road (Nvertspecies~Distance; p=0.22, R^2^=0.18). Thus, overall, there is no evidence that vertebrate species richness is dependent on sampling flies far from the road. However, the examination of Shannon diversity for vertebrates across the different sampling sites and distance from disturbance suggests that less sampling is needed within the forest in order to detect most of the vertebrates (Figure 1D).

**Figure 3.**
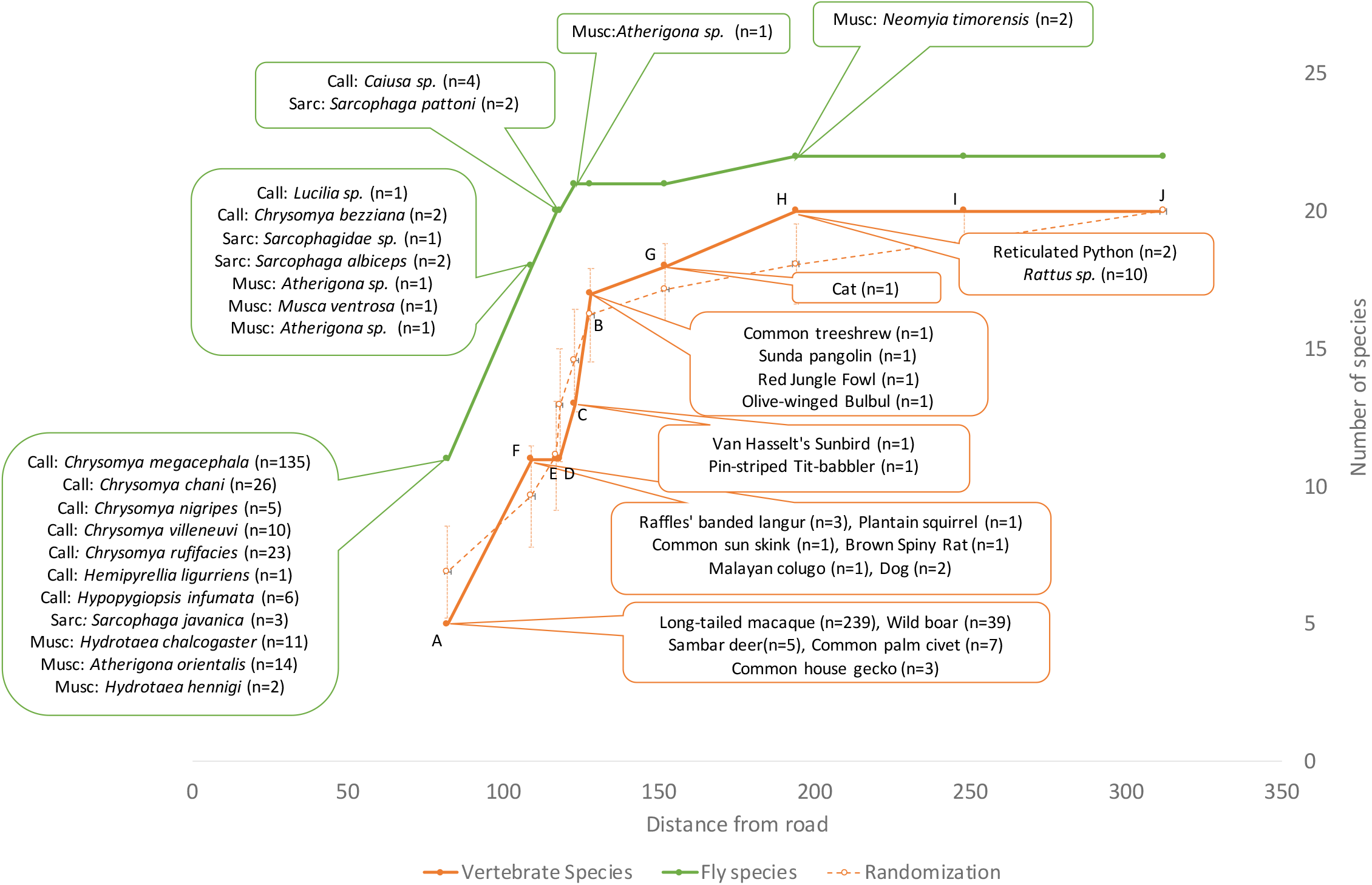
Identity and cumulative number of vertebrates (orange) and flies (green) identified as sampling is carried out at sites deeper into the forest. Dashed line shows the predicted vertebrate diversity obtained when flies are randomized across sites. Total number of barcoded flies=255, flies without barcodes containing vertebrate hits: 15.

### Factors determining vertebrate detection in a fly sample

We assessed the probability of detecting a vertebrate (irrespective of the species being a free-ranging animal or not) as a function of the sex of the fly, its family membership, or the type of sample (e.g., feces, regurgitate). Logistic regression models suggests that the probability of detecting vertebrate species is not well explained by the sex of the fly (Supplementary Table 2). On the other hand, the models using family membership or sample type show that there is a significant association between these variables and the detection of vertebrate species. However, given that these variables may interact, we implemented a model accounting for the interactions. This model show that sample type is significantly associated with failure to detect vertebrate species. Note however that, a visualization of detection of vertebrate species vs the type of sample suggests that all types of material can yield vertebrate identifications (Supplementary Figure 1).

### Towards MinION based vertebrate surveillance with iDNA

Lastly, we tested whether future studies could use MinION sequencing. Overall, we obtained 6,027,547 nanopore sequences for 313-bp COI fragment, 5,020,129 for 244-bp COI fragment and 7,312,601 for 16S. Demultiplexing of these data yielded 17.51%, 39.67% and 42.03% of reads binned to specimens. A total of 68, 20, and 330 flies gave matches to vertebrates for 313-bp COI, 244-bp COI and 16S fragments respectively at 95% distance threshold. When this threshold was lowered to 90%, the corresponding numbers were 73, 23 and 312. The reduction in number of matches with lower identity criterion for 16S was due to do the higher read count threshold established due to greater number of sequences in negative (8 vs 11 for 95% vs 90%). Compared to 95% identity criterion, the application of a 90% identity criterion yielded fewer matches, more conflict with identifications based on Illumina (Supplementary Table 3, Summary statistics), and more unexpected matches (e.g. presence of Silvery gibbon *Hylobates moloch* and larger number of reads matching Rhesus macaque *Macaca mulatta* and Chimpanzee *Pan troglodytes:* Supplementary Table 3). Therefore, subsequent analyses only used the 95% identity criterion.

The combined vertebrate matches based on all 3 markers for Illumina and MinION yield very similar species interaction networks (Figure 4 A,B). Of the six unusual identifications (yellow, red in Figure 4 A,B), three are due to greater amount of ambiguity in MinION based identification (*Gallus sp*. instead of *Gallus gallus*, or *Cervidae sp*. instead of *Rusa unicolor*). However, MinION reads suggest the presence of *Trachypithecus fransoici* for a sample that contains old-world monkey (*Presbystis femoralis*) based on Illumina reads. Similarly, *Anser canagicus* co-occurred with *Gallus sp*. in MinION data. The last unusual identification was *Pan troglodytes*, which likely co-occurred with *Homo sapiens* that was filtered out of our datasets. Some issues are found for both Illumina and MinION datasets (e.g., multiple *Macaca mulatta* identifications). These are largely eliminated in our full analysis of the Illumina data using multiple replicates and/or translation check. Nevertheless, these discrepancies are minor compared to the >300 identifications that are comparable across the datasets (Supplementary Tables 3,5). When the effect of disturbance gradient was explored, MinION data revealed that roadside sampling is sufficient, and paralleled the plot obtained by Illumina data. However, the species may be accumulated at different points along the gradient, largely due to varying coverages and filtering criteria (Figure 5).

**Figure 4.**
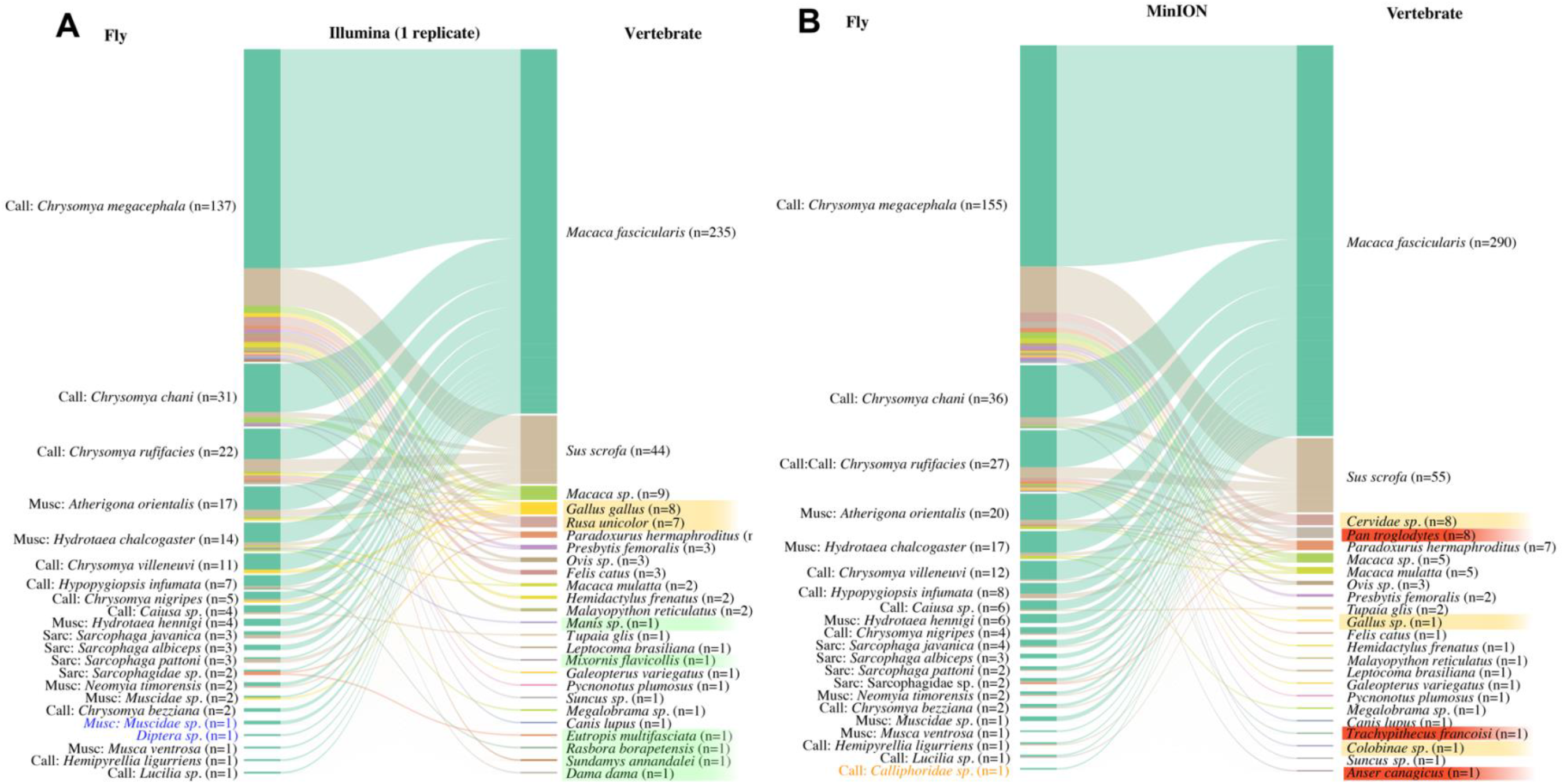
Species interaction network based on (A) Illumina reads and (B) MinION reads (95% identity) using one PCR replicate. Text color: blue=Illumina only, orange=MinION only. Background color: green=likely accurate identification found only in one dataset; red=likely inaccurate identification found only in one dataset; yellow= identification to genus/family based on MinION data but species specific for Illumina.

**Figure 5.**
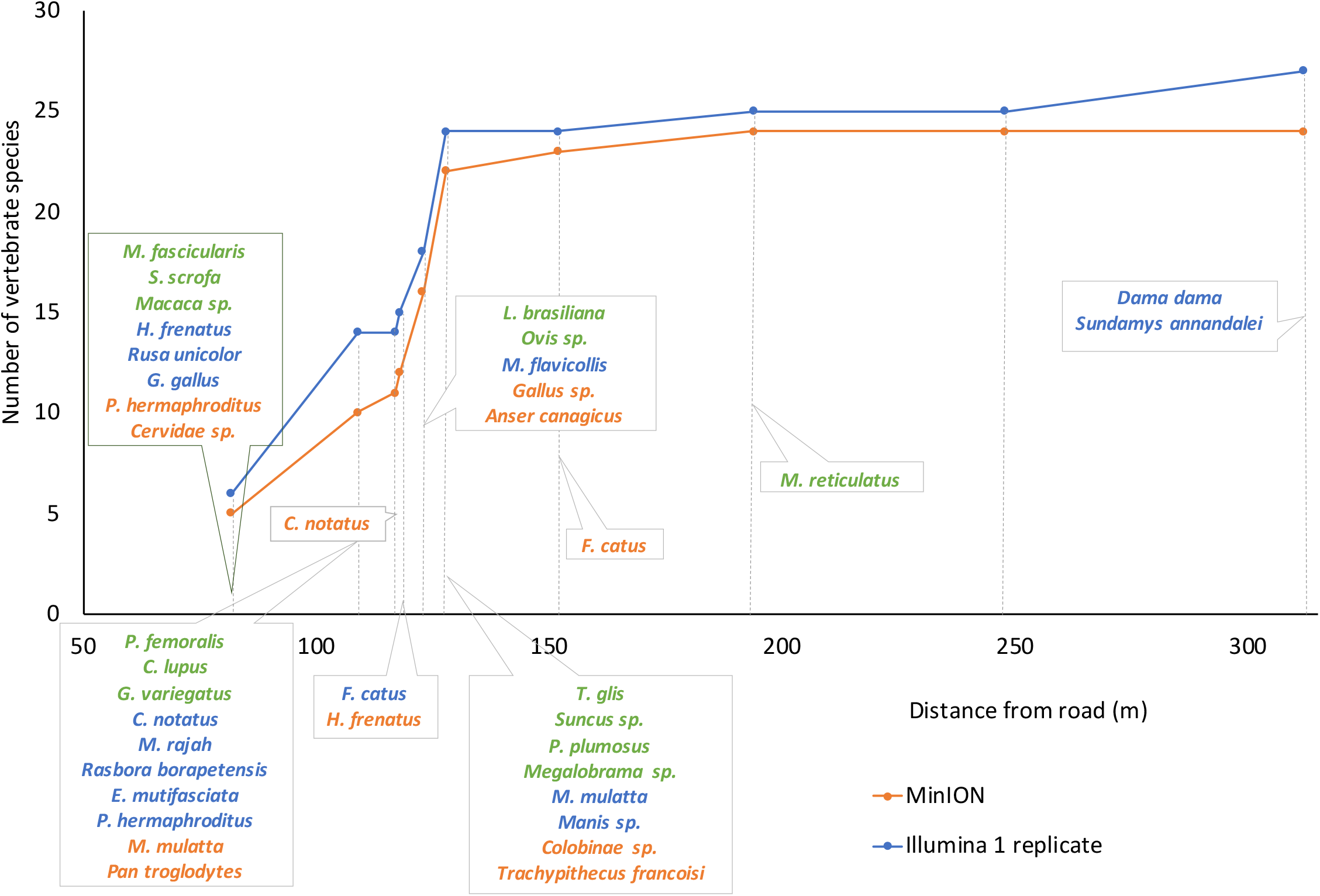
Vertebrate detection curve for MinION and Illumina datasets. This technical comparison includes matches to non-free-ranging animals and unusual identifications. Green=both platforms, blue=Illumina only, orange=MinION only.

## DISCUSSION

We here tested whether iDNA from flies attracted to carrion in a swamp forest show signs of specialization that would allow for optimizing sampling strategies and providing new insights into the ecology of the forest. We did not find significant specialization in a quantitative analysis of the fly-vertebrate interaction, but the sample sizes for sarcophagid and muscid flies were too small to draw robust conclusions. Our study does, however, show that even rare vertebrate species can be detected based on flies collected within 150m of a road that dissects a high conservation-value swamp forest remnant that houses many endangered vertebrate species. iDNA was effective at detecting rare ground-dwelling and arboreal species such as the Sunda pangolin (*Manis javanica*) and Raffles’ banded langur (*Presbytis femoralis*) and also confirmed high abundance of species like long-tailed macaque (*Macaca fascicularis*) and wild boar (*Sus scrofa*) that are routinely sighted in the field site. In addition, we show that all the data in our study can be obtained using a greatly simplified iDNA sampling and processing protocol that avoids DNA extraction. The overall results are furthermore consistent between Illumina and Nanopore sequencing.

A major challenge in monitoring forest vertebrates is the logistics of field surveys, especially in forests with limited accessibility. We here show that iDNA obtained from flies baited near a road (~150 m) contained DNA for almost all vertebrate species in our study (18 of 20; Figure 3, see also linear model). Additional sampling deeper in the forest only revealed two additional species that are so common in urban environments that their detection is of little significance (reticulated python and Asian house/brown rat). Note that the fly community close to the road was more diverse than the community within the forest. This included some species that were less likely to yield vertebrate records, but contained signal for vertebrate species rarely encountered in our study. We would thus recommend that dense roadside sampling is preferable to only sampling flies within the forest because the latter is more time-consuming and invasive.

### Lack of specialization in fly-vertebrate interactions

We sampled approximately 400 flies that belong to 25 species within Calliphoridae, Muscidae, and Sarcophagidae. Most vertebrate records came from only three calliphorid species belonging to *Chrysomya* (219 records for 14 species). The main “workhorse” of our and other iDNA studies in Southeast Asia was *Chrysomya megacephala* (Figure 2, Lee et al., 2016, 2015); it contributes 163 records for 13 species based on 135 flies. Our results nevertheless suggest that all *Chrysomya* specimens should be analyzed because they yield identifications for mammals, birds and reptiles. Most *Chrysomya* contained strong signal for long-tailed macaque and wild boar and we would thus advise against pooling because the signal for rarer vertebrate species may be overlooked if the flies were not analyzed individually (Lynggaard et al., 2019).

Overall, our network-level analysis for all specimens and records failed to reveal a significant levels of specialization (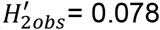, P=0.712 for species level, 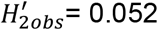, P=0.33 for family level), but this could be due to sampling given that we had only fairly few specimens for 23 of the 25 flies species. Indeed, based on a very small sample size (N=7), we find limited evidence for greater vertebrate specialization in fly species belonging to Sarcophagidae. Muscid flies had lower Shannon diversity for vertebrates compared to sarcophagids and calliphorids but yielded the only record for the endangered Sunda pangolin. Indeed, excluding all muscid and sarcophagid species from analysis would lead to the loss of several vertebrate species (*Manis*, *Eutropis*) and the number of records for some other species would be reduced (e.g., *Paradoxurus*). Only analyzing the “classic” *Chrysomya* blow flies would eliminate another two vertebrate species from the list of detected species. These qualitative observations imply that it would be important in future studies to sample more muscids and sarcophagids which is realistic if different baits were used (e.g., mammal dung). Such baits would also attract insects from other orders (e.g., beetles: Drinkwater et al., 2021). Alternative artificial/chemical baits and commercial traps should be further explored for easier sampling. All work on these iDNA carriers should include specimen-level barcoding in order to understand species interactions. Such barcoding is now very simple and cost-effective (Meier, Wong, Srivathsan, & Foo, 2016; Srivathsan et al., 2021; Yeo et al., 2021).

More detailed work on the natural history of flies is also important for addressing one of the least satisfying aspects of iDNA studies. We only have limited information about how far invertebrates can disperse. Carrion flies may move 100–2400 m per (Lee et al., 2016) which could explain how a *Chrysomya megacephala* contained iDNA signal from a kangaroo (kangaroos are found in the Singapore Zoo). Such long-distance dispersal was previously invoked to explain *Canis* signal on Barro Colorado Island in Panama (Rodgers et al., 2017). In this context, it is worth noting that blow fly guts can contain amplifiable DNA 24-96 hours post feeding (Lee et al., 2015).

The most commonly detected vertebrate species in our survey were long-tailed macaques (*Macaca fascicularis*), and wild boars (*Sus scrofa*), but we also detected rare species. Particularly noteworthy were two hits for critically endangered species. Two hits were for Raffles’ banded langur (*Presbytis femoralis*), which is a globally Critically Endangered species with ~60 individuals remaining in Singapore and overall global population of <250 mature individuals (Ang et al., 2020). Field observations of this species are difficult because it is shy and elusive (Ang et al., 2010). Its detection with iDNA confirms that flies are an excellent tool for monitoring rare mammals including arboreal primates that are difficult to monitor with camera traps (see Gogarten et al., 2020). Similarly remarkable is the detection of a Sunda pangolin (*Manis javanica*), which is a nationally critically endangered species with a population estimate of only ~1000 individuals based on camera trap surveys, radio telemetry, and mark-and-recapture information from rescued pangolins. Data acquisition for this species has been extremely difficult (Nash et al., 2020) and iDNA will be a welcome additional tool. In addition to critically endangered species, we also found evidence for one near-threatened species: the Malayan colugo (*Galeopterus variegatus*), which is a forest-dwelling and one vulnerable species: the Brown spiny rat (*Maxomys rajah*), which is usually found only in primary and secondary forests (Nakagawa et al., 2009). Finding such rare and endangered species is not unusual in iDNA studies. Leech data have previously revealed critically endangered and even newly discovered mammal species in Vietnam (Schnell et al., 2012).

In addition to bird taxa that are unlikely to be detected by camera traps, we also find evidence for the presence of several small-bodied species that are either unidentifiable on camera trap images or unlikely to be photographed (e.g., common house gecko, common sun skink, Asian rat, common treeshrew). Detecting some of these species would have traditionally required the use of invasive trapping methods. We thus used species richness estimation methods to establish how many additional vertebrate species could be obtained with additional sampling (Figure 1). It suggests that another 30 species is a realistic estimate. This is still below the known vertebrate species diversity of Nee Soon and highlights once more that complete vertebrate surveys require the use of multiple tools. One potential way to increase the proportion of rare species detected would be the use of blocking primers for the dominant *Macaca fascicularis* and *Sus scrofa*. However, blocking primers for a common primate may also interfere with the detection of a critically endangered one (*Presbytis femoralis*).

### A simplified procedure for iDNA based vertebrate monitoring

Our results suggest that a greatly simplified iDNA protocol can efficiently detect vertebrates. DNA template is simply obtained by dissolving fly feces and regurgitates with water, followed by PCR and sequencing. Using this protocol, >60% of the flies generated >300 vertebrate detections and revealed a substantial proportion of the known vertebrate diversity of the study site (Figure 1). It is remarkable that this detection frequency is higher than the detection rate of vertebrates in experimental studies using DNA extracted from whole flies (Calvignac-Spencer et al. (2013): 39%, Lee et al (2015): 27%). This suggests that our simplified protocol is highly effective and suitable for the rapid surveillance of vertebrate diversity. This is welcome news given that understanding of vertebrate communities with iDNA would require large-scale sampling and the processing of hundreds or thousands of flies; i.e., the molecular methods need to be simple and cost-effective. In many ways, our simplified molecular protocol is matched by a similarly simple sampling protocol that only requires a foldable butterfly cage, bait, and sterile plastic tubes. The latter helps with avoiding contamination, because once a fly has been caught in the vial, it does not have to be opened again until it reaches the lab. This makes our protocol also suitable for use by citizen scientists. We also tested whether feces, regurgitates, or seemingly empty vials are more reliable sources of iDNA signal. A linear model detected that the sample type “Empty” and “Clear” had lower probability of detections compared to samples containing both “Brown” and “Clear” material (Clear: p=0.001, Empty: p=0.0004) (Supplementary Table 2). We suggest that more research is needed to understand whether the different sources of iDNA should be analyzed separately. This may be desirable, because they may reflect when the vertebrate iDNA was acquired. It is conceivable that “empty” vials only contain very fresh vertebrate DNA transferred via fly legs. Regurgitates, on the other hand, would be somewhat older if they came from the long crop of flies. Lastly, fly feces should contain the “oldest” evidence because it consists of digested food.

The workflow proposed in our study has three major benefits over traditional approaches. Firstly, it is simple to obtain the samples and the flies do not have to be handled. Yet, the vast majority of the flies in this study produced feces, regurgitates, or both (92%). In previous studies involving flies, blood meals either had to be squeezed out of the gut or the entire specimen had to be lysed. Secondly, DNA extraction is skipped and there is no need for kits or buffers (Calvignac-Spencer et al., 2013; Reeves et al., 2018). This greatly reduces cost, time, and labor. Thirdly, our metabarcoding experiments use a tagging strategy that only requires a single PCR with tagged primers. Chimera formation is avoided by employing PCR-free DNA libraries where adapters are ligated to amplicons. Currently, most studies still use two rounds of PCR for indexing samples (Lee et al., 2016; Rodgers et al., 2017), which increases the chemical cost, time and labor involved in the experiments (see comments in Srivathsan et al. 2021).

Lastly, we demonstrate that Nanopore sequencing is now sufficiently accurate to yield results that are broadly congruent with what is found with Illumina sequencing. MinION still gives a few additional erroneous matches (e.g., *T. fransoici*) but these could be eliminated based on a checklist for the site and country. Overall, our results suggests that MinION is quickly becoming a powerful tool for biodiversity surveys using metabarcoding. There are constant improvements in MinION sequencing chemistry and base-calling. With the use of recent kits and new base-calling algorithms, we were able to identify a large number of vertebrate species at a 95% distance threshold. The amount of data that can be obtained per MinION flowcell now depends largely on primer specificity. For example, with the 313-bp COI fragment, which co-amplifies host DNA, the amount of vertebrate signal was lower, and we estimate that 100 vertebrate samples could be multiplexed in a run (sample cost of ~10 USD), but for 16S we were able to get identifications from >300 samples in one run. We nevertheless estimate that analysis with MinION is still one order of magnitude more expensive than what would be needed for Illumina; i.e. large-scale studies should continue to use this platform because it also facilitates the analysis of multiple PCR replicates.

The effectiveness of our protocols can be further improved by choosing more and/or different primer pairs. Each of the primer pairs used here had its advantages and limitations. The COI minibarcode amplifying 313-bp fragment co-amplified host and gave few reads for vertebrate identifications. This primer pair was able to amplify mammals and reptiles. However, analysis of the data with just this pair yielded a species interaction network that included some records that were not present in the combined dataset based on 16S and a different section of COI. The second COI minibarcode was problematic in that it co-amplified bacteria and yielded the lowest number of flies with vertebrate identifications (N=34). However, this primer was effective in detecting of birds. Overall, the vast majority of vertebrate detections (N=303) were based on 16S, but they only pertained to mammals. These findings suggest that it is critical to combine multiple markers in iDNA surveys.

## CONCLUSION

iDNA has repeatedly proven to be a useful tool for surveying vertebrate diversity. However, the experimental costs and labor involved have been major deterrents to widespread adoption, particularly in biodiverse region. In this study, we develop a protocol that greatly simplifies the molecular procedures and shows that nanopore sequencing can allow for on-site data generation. We demonstrate that sampling close to a road is sufficient for obtaining almost all vertebrate hits. However, when it comes to interpreting the results, it will be important to know more about the biology of iDNA sources and to study species interactions based on larger sample sizes.

## ACKNOWLEDGEMENTS

We would like to thank the National Parks board for allowing us bait and collect the fly samples. We would also like to thank Glendon Kee and Amos Chua for their assistance with the fly sampling.

## DATA ACCESSIBILITY

All Illumina and MinION fastq files, related demultiplexing information and the reference barcodes are available in Dryad (Srivathsan, 2022): https://doi.org/10.5061/dryad.2547d7wtj. Script for randomization is available in GitHub: https://github.com/asrivathsan/iDNA_roadside.

## AUTHOR CONTRIBUTIONS

EO: sampling and initial experimentation. RLK and SNK: molecular work, nanopore sequencing, and analysis AS: analysis, writing and experimental optimizations, LL: DNA barcoding, YA: fly morphology, RM: conception, guidance and writing.

