## Supplementary for "Network analysis with either Illumina or MinION reveals that detecting vertebrate species requires metabarcoding of iDNA from a diverse fly community"

**Supplementary Table 1:** Metadata for sampling units. Data for barcoded flies only.

| **Sampling point** | **Coordinate** | **Distance from road** | **Number of flies collected** | **Number of fly species** | **Number of flies with vertebrates** | **Number of fly species with vertebrates** |
| --- | --- | --- | --- | --- | --- | --- |
| A | 1.3816617, 103.8154960 | 82 | 71 | 14 | 45 | 11 |
| B | 1.3819885, 103.8157348 | 128 | 73 | 11 | 32 | 7 |
| C | 1.3822711, 103.8153301 | 123 | 39 | 12 | 26 | 10 |
| D | 1.3824822, 103.8148221 | 118 | 37 | 8 | 24 | 7 |
| E | 1.382368, 103.815032 | 117 | 29 | 12 | 18 | 8 |
| F | 1.3824933, 103.8144989 | 109 | 48 | 15 | 33 | 13 |
| G | 1.3824299, 103.8155638 | 152 | 26 | 3 | 15 | 2 |
| H | 1.3828643, 103.8155933 | 194 | 28 | 4 | 21 | 4 |
| I | 1.3832833, 103.8158327 | 248 | 26 | 4 | 17 | 2 |
| J | 1.3838404, 103.8160083 | 312 | 30 | 4 | 22 | 3 |

Supplementary Table 2: Summary statistics for logistic regression model for detection of vertebrate species

| **Model** | **Reference** | **Category assessed** | **Coefficient** | **SE** | **P** |
| --- | --- | --- | --- | --- | --- |
| vertpresence~sex | Female | Male | -0.2876 | 0.2534 | 0.256 |
| vertpresence~family | Calliphoridae | Muscidae | -1.1027 | 0.2704 | 4.53e-05 *** |
| vertpresence~family | Calliphoridae | Sarcophagidae | -0.5782 | 0.5695 | 0.31 |
| vertpresence~sample.type | Brown + Clear | Brown | -0.1985 | 0.3612 | 0.582695 |
| vertpresence~sample.type | Brown + Clear | Clear | -0.7887 | 0.2417 | 0.001101** |
| vertpresence~sample.type | Brown + Clear | Empty | -1.4464 | 0.4116 | 0.000442*** |

**Supplementary Table 3:** Comparison of identifications made by Illumina and MinION metabarcoding experiments for the 3 genes and different identity cut-offs for MinION. Values in Species section reflect number of flies with identifications made by Illumina/MinION (90% cutoff)/MinION (95% cut-off). Summary Statistics reflect comparisons of MinION (90%) vs Illumina/MInION (95%) vs illumina)

| **Species** | **313 bp COI (I/M90/M95)** | **244 bp COI**  **(I/M90/M95)** | **16S (I/M90/M95)** |
| --- | --- | --- | --- |
| *Anser canagicus* | 0/0/0 | 0/1/1 | 0/0/0 |
| *Callosciurus notatus* | 0/0/0 | 0/0/0 | 2/1/1 |
| *Canis lupus* | 1/0/0 | 0/0/0 | 1/1/1 |
| *Cervidae sp.* | 0/0/0 | 0/0/0 | 0/8/8 |
| *Colobinae sp.* | 0/0/0 | 0/0/0 | 0/1/1 |
| *Dama dama* | 0/0/0 | 1/0/0 | 0/0/0 |
| *Eutropis multifasciata* | 0/0/0 | 1/0/0 | 0/0/0 |
| *Felis catus* | 2/0/0 | 0/0/0 | 1/1/1 |
| *Galeopterus variegatus* | 0/0/0 | 0/0/0 | 2/2/2 |
| *Gallus gallus* | 0/0/0 | 8/0/0 | 0/0/0 |
| *Gallus sp.* | 0/0/0 | 0/1/1 | 0/0/0 |
| *Hemidactylus frenatus* | 0/0/0 | 3/1/1 | 0/0/0 |
| *Hylobates moloch* | 0/0/0 | 0/0/0 | 0/1/0 |
| *Leptocoma brasiliana* | 0/0/0 | 1/1/1 | 0/0/0 |
| *Macaca fascicularis* | 75/59/57 | 9/5/5 | 257/292/312 |
| *Macaca mulatta* | 2/3/2 | 3/3/1 | 3/7/6 |
| *Macaca nemestrina* | 2/2/2 | 0/0/0 | 0/0/0 |
| *Macaca sp.* | 0/0/0 | 3/0/0 | 0/0/0 |
| *Malayopython reticulatus* | 1/0/0 | 2/1/1 | 0/0/0 |
| *Manis sp.* | 0/0/0 | 0/0/0 | 1/0/0 |
| *Maxomys rajah* | 0/0/0 | 1/0/0 | 0/0/0 |
| *Megalobrama sp.* | 1/1/1 | 0/0/0 | 0/0/0 |
| *Mixornis flavicollis* | 1/0/0 | 0/0/0 | 0/0/0 |
| *Mus pahari* | 0/0/0 | 0/0/0 | 0/5/0 |
| *Ovis aries* | 1/1/1 | 0/0/0 | 0/0/0 |
| *Ovis sp.* | 0/0/0 | 0/0/0 | 3/3/3 |
| *Pan troglodytes* | 0/0/0 | 0/0/0 | 0/9/8 |
| *Paradoxurus hermaphroditus* | 2/2/1 | 1/1/1 | 5/4/6 |
| *Presbytis femoralis* | 2/1/1 | 0/0/0 | 3/2/2 |
| *Pycnonotus plumosus* | 1/1/1 | 0/0/0 | 0/0/0 |
| *Rasbora borapetensis* | 0/0/0 | 0/0/0 | 1/0/0 |
| *Rusa unicolor* | 0/0/0 | 2/1/1 | 6/0/0 |
| *Suncus sp.* | 0/0/0 | 0/0/0 | 1/1/1 |
| *Sundamys annandalei* | 0/0/0 | 1/0/0 | 0/0/0 |
| *Sus scrofa* | 9/3/3 | 28/11/10 | 29/42/57 |
| *Trachypithecus francoisi* | 0/0/0 | 0/0/0 | 0/1/1 |
| *Tupaia glis* | 1/1/1 | 0/0/0 | 0/2/2 |
| **Summary Statistics** | **313 bp COI  (I vs M90/ I vs M95)** | **244 bp COI**  **(I vs M90/ I vs M95)** | **16S  (I vs M90/ I vs M95)** |
| Present in both | 71/69 | 21/19 | 285/292 |
| MinION only, present at lower coverage cutoff in Illumina | 0/0 | 0/0 | 69/99 |
| Minion only, species specific match in Illumina | 0/0 | 3/3 | 7/7 |
| Minion only, in Illumina at lower identity cutoff | 2/1 | 1/0 | 14/11 |
| Minion only | 1/0 | 1/1 | 8/3 |
| Illumina Only | 33/35 | 43/45 | 23/23 |

**Supplementary Table 4: Percentage identities for vertebrate matches for the analysis of iDNA metabarcoding using Illumina (2 replicates)**

| **Sample** | **Vertebrate** | **313 bp COI** | **244 bp COI** | **16S** |
| --- | --- | --- | --- | --- |
| A002 | Sus scrofa | NA | 100 | 100 |
| A009 | Sus scrofa | NA | 100 | NA |
| A015 | Sus scrofa | NA | 100 | NA |
| A017 | Rusa unicolor | NA | NA | 100 |
| B078 | Macaca fascicularis | 100 | NA | 100 |
| B080 | Macaca fascicularis | NA | NA | 100 |
| B081 | Presbytis femoralis | NA | NA | 100 |
| B082 | Macaca fascicularis | NA | NA | 100 |
| B084 | Macaca fascicularis | NA | NA | 100 |
| B085 | Macaca fascicularis | 100 | NA | 100 |
| B086 | Macaca fascicularis | NA | NA | 100 |
| B088 | Macaca fascicularis | NA | NA | 100 |
| B089 | Macaca fascicularis | NA | NA | 100 |
| B093 | Macaca fascicularis | NA | NA | 100 |
| B095 | Macaca fascicularis | NA | NA | 100 |
| B095 | Manis javanica,Manis pentadactyla | NA | NA | 100 |
| C153 | Sus scrofa | NA | 100 | NA |
| C156 | Macaca fascicularis | NA | NA | 100 |
| C157 | Macaca fascicularis | NA | NA | 100 |
| C158 | Macaca fascicularis | NA | NA | 100 |
| C160 | Macaca fascicularis | 100 | NA | 100 |
| C161 | Macaca fascicularis | NA | NA | 100 |
| C162 | Macaca fascicularis | NA | NA | 100 |
| C163 | Macaca fascicularis | NA | NA | 100 |
| C164 | Macaca fascicularis | 99.681 | NA | 100 |
| D197 | Macaca fascicularis | 99.681 | NA | NA |
| D197 | Sus scrofa | NA | 100 | NA |
| D198 | Macaca fascicularis | 100 | NA | 100 |
| D199 | Sus scrofa | NA | 100 | NA |
| D200 | Macaca fascicularis | 100 | NA | NA |
| D201 | Bos taurus | NA | NA | 100 |
| D201 | Rusa unicolor | NA | NA | 100 |
| D204 | Paradoxurus hermaphroditus | NA | NA | 100 |
| A020 | Macaca fascicularis | NA | NA | 100 |
| A022 | Macaca fascicularis | NA | NA | 100 |
| A024 | Macaca fascicularis | NA | NA | 100 |
| A025 | Macaca fascicularis | NA | NA | 100 |
| A026 | Macaca fascicularis | 99.681 | NA | 100 |
| A027 | Macaca fascicularis | NA | NA | 100 |
| A028 | Macaca fascicularis | 99.681 | NA | 100 |
| A029 | Macaca fascicularis | NA | NA | 100 |
| A030 | Macaca fascicularis | NA | NA | 100 |
| A031 | Macaca fascicularis | NA | NA | 100 |
| A032 | Macaca fascicularis | NA | NA | 100 |
| A035 | Macaca fascicularis | NA | NA | 100 |
| A036 | Macaca fascicularis | 99.681 | NA | 100 |
| A037 | Macaca fascicularis | NA | NA | 100 |
| A039 | Macaca fascicularis | NA | NA | 100 |
| A040 | Macaca fascicularis | 100 | NA | 100 |
| A041 | Macaca fascicularis | NA | NA | 100 |
| A042 | Macaca fascicularis | NA | NA | 100 |
| A043 | Macaca fascicularis | NA | NA | 100 |
| A044 | Macaca fascicularis | NA | NA | 100 |
| A045 | Macaca fascicularis | 99.681 | NA | 100 |
| A046 | Macaca fascicularis | NA | NA | 100 |
| A047 | Macaca fascicularis | NA | NA | 100 |
| B098 | Macaca fascicularis | NA | NA | 100 |
| B099 | Macaca fascicularis | NA | NA | 100 |
| B100 | Macaca fascicularis | NA | NA | 100 |
| B100 | Sus scrofa | NA | NA | 100 |
| B103 | Sus scrofa | NA | 100 | 100 |
| B103 | Macaca fascicularis | NA | NA | 100 |
| B104 | Tupaia glis | 99.681 | NA | NA |
| B104 | Macaca fascicularis | NA | NA | 100 |
| B105 | Macaca fascicularis | 99.681 | NA | 100 |
| B106 | Macaca fascicularis | NA | NA | 100 |
| B108 | Macaca mulatta | NA | NA | 98.936 |
| B111 | Macaca fascicularis | NA | NA | 100 |
| B115 | Macaca fascicularis | NA | NA | 100 |
| B118 | Macaca fascicularis | NA | NA | 100 |
| B118 | Sus scrofa | NA | NA | 100 |
| B119 | Macaca fascicularis | NA | NA | 100 |
| B121 | Gallus gallus | NA | 100 | NA |
| B122 | Sus scrofa | 100 | NA | 100 |
| B122 | Macaca fascicularis | NA | NA | 100 |
| C167 | Macaca fascicularis | 99.681 | NA | 100 |
| C169 | Macaca fascicularis | 99.681 | NA | 100 |
| C170 | Macaca fascicularis | NA | NA | 100 |
| C173 | Macaca fascicularis | 99.681 | NA | 100 |
| C173 | Leptocoma brasiliana | NA | 99.083 | NA |
| C174 | Gallus gallus | NA | 100 | NA |
| C176 | Macaca fascicularis | 100 | NA | 100 |
| C176 | Canis lupus | NA | NA | 100 |
| C177 | Macaca fascicularis | NA | NA | 100 |
| C178 | Macaca fascicularis | 99.681 | NA | 100 |
| C178 | Mixornis flavicollis | 95.793 | NA | NA |
| C181 | Macaca fascicularis | NA | NA | 100 |
| C181 | Ovis sp. | NA | NA | 100 |
| C182 | Macaca fascicularis | 99.681 | NA | 100 |
| C183 | Macaca fascicularis | NA | NA | 100 |
| C184 | Macaca fascicularis | 100 | NA | 100 |
| C187 | Ovis sp. | NA | NA | 100 |
| C188 | Macaca fascicularis | 99.681 | NA | 100 |
| C190 | Macaca fascicularis | 99.681 | NA | 100 |
| C191 | Hemidactylus frenatus | NA | 100 | NA |
| C191 | Macaca fascicularis | NA | NA | 100 |
| C192 | Macaca fascicularis | 99.681 | NA | 100 |
| C193 | Macaca fascicularis | 99.681 | NA | 100 |
| C194 | Macaca fascicularis | NA | NA | 100 |
| D207 | Macaca fascicularis | NA | NA | 100 |
| D208 | Macaca fascicularis | NA | NA | 100 |
| D209 | Macaca fascicularis | NA | NA | 100 |
| D210 | Presbytis femoralis | 100 | NA | 100 |
| D214 | Macaca fascicularis | 100 | NA | 100 |
| D215 | Macaca fascicularis | NA | NA | 100 |
| D216 | Macaca fascicularis | 99.681 | 100 | 100 |
| D216 | Hemidactylus frenatus | NA | 100 | NA |
| D220 | Macaca fascicularis | NA | NA | 100 |
| D221 | Macaca fascicularis | NA | NA | 100 |
| D224 | Paradoxurus hermaphroditus | 96.711 | NA | NA |
| D224 | Macaca fascicularis | NA | NA | 100 |
| D225 | Macaca fascicularis | NA | NA | 100 |
| D226 | Macaca fascicularis | NA | NA | 100 |
| D227 | Sus scrofa | 100 | NA | NA |
| D227 | Macaca fascicularis | NA | NA | 100 |
| D228 | Macaca fascicularis | 100 | NA | 100 |
| D229 | Macaca fascicularis | NA | NA | 100 |
| D230 | Macaca fascicularis | NA | NA | 100 |
| D232 | Sus scrofa | NA | 100 | NA |
| D233 | Macaca fascicularis | NA | NA | 100 |
| D235 | Macaca fascicularis | NA | NA | 100 |
| D236 | Macaca fascicularis | NA | NA | 100 |
| A048 | Hemidactylus frenatus | NA | 100 | NA |
| A048 | Macaca fascicularis | NA | NA | 100 |
| A049 | Macaca fascicularis | 100 | NA | 100 |
| A050 | Macaca fascicularis | NA | NA | 100 |
| A051 | Macaca fascicularis | NA | NA | 100 |
| A052 | Macaca fascicularis | NA | NA | 100 |
| A054 | Macaca fascicularis | NA | NA | 100 |
| A055 | Macaca fascicularis | NA | NA | 100 |
| A056 | Paradoxurus hermaphroditus | NA | NA | 100 |
| A057 | Macaca fascicularis | NA | NA | 100 |
| A058 | Macaca fascicularis | NA | NA | 100 |
| A060 | Macaca fascicularis | NA | NA | 100 |
| A061 | Macaca fascicularis | NA | NA | 100 |
| A063 | Macaca fascicularis | NA | NA | 100 |
| A064 | Macaca fascicularis | NA | NA | 100 |
| A067 | Macaca fascicularis | 99.681 | NA | 100 |
| A068 | Macaca fascicularis | 100 | NA | 100 |
| B128 | Sus scrofa | NA | 100 | NA |
| B130 | Sus scrofa | NA | NA | 100 |
| B134 | Sus scrofa | NA | 100 | 100 |
| B135 | Macaca fascicularis | 100 | 100 | NA |
| B135 | Pycnonotus plumosus | 99.042 | NA | NA |
| B137 | Sus scrofa | NA | NA | 100 |
| B141 | Macaca sp. | NA | 100 | NA |
| B141 | Sus scrofa | NA | 100 | NA |
| B141 | Macaca mulatta | NA | NA | 100 |
| E237 | Macaca fascicularis | NA | NA | 100 |
| E239 | Macaca fascicularis | NA | NA | 100 |
| E240 | Macaca fascicularis | NA | NA | 100 |
| E242 | Macaca fascicularis | NA | NA | 100 |
| E243 | Macaca fascicularis | 99.681 | NA | 100 |
| E244 | Macaca fascicularis | NA | NA | 100 |
| E245 | Macaca fascicularis | NA | NA | 100 |
| E246 | Macaca fascicularis | NA | NA | 100 |
| E248 | Macaca fascicularis | NA | NA | 100 |
| E250 | Sus scrofa | NA | NA | 100 |
| E251 | Macaca fascicularis | 99.681 | 100 | 100 |
| E253 | Macaca fascicularis | NA | NA | 100 |
| E254 | Macaca fascicularis | NA | NA | 100 |
| E257 | Macaca sp. | NA | 100 | NA |
| E257 | Macaca mulatta | NA | NA | 100 |
| F270 | Macaca fascicularis | NA | NA | 100 |
| F271 | Macaca fascicularis | NA | NA | 100 |
| F272 | Macaca fascicularis | NA | NA | 100 |
| F273 | Macaca fascicularis | NA | NA | 100 |
| F274 | Macaca fascicularis | NA | NA | 100 |
| F275 | Macaca fascicularis | NA | NA | 100 |
| F276 | Macaca fascicularis | NA | NA | 100 |
| F277 | Macaca fascicularis | NA | NA | 100 |
| F279 | Sus scrofa | NA | 100 | NA |
| F280 | Rusa unicolor | NA | 100 | 100 |
| F280 | Sus scrofa | NA | 100 | 100 |
| F280 | Macaca fascicularis | NA | NA | 100 |
| F285 | Macaca fascicularis | NA | NA | 100 |
| F289 | Macaca fascicularis | 100 | NA | 100 |
| F290 | Macaca fascicularis | NA | NA | 100 |
| A073 | Macaca fascicularis | NA | NA | 100 |
| A074 | Macaca fascicularis | NA | NA | 100 |
| A075 | Macaca fascicularis | NA | NA | 100 |
| A077 | Macaca fascicularis | 100 | NA | 100 |
| B147 | Macaca fascicularis | 100 | NA | 100 |
| B148 | Macaca fascicularis | NA | NA | 100 |
| B151 | Macaca fascicularis | NA | NA | 100 |
| E259 | Macaca fascicularis | 100 | NA | 100 |
| E260 | Macaca fascicularis | NA | NA | 100 |
| E261 | Macaca fascicularis | 99.681 | NA | 100 |
| E261 | Sus scrofa | 100 | 100 | 100 |
| E263 | Macaca fascicularis | NA | NA | 100 |
| E265 | Sus scrofa | NA | 100 | 100 |
| E266 | Macaca fascicularis | 100 | 100 | 100 |
| E267 | Sus scrofa | NA | 100 | 100 |
| E268 | Macaca fascicularis | NA | NA | 100 |
| F291 | Macaca fascicularis | NA | NA | 100 |
| F292 | Presbytis femoralis | 100 | NA | 100 |
| F292 | Sus scrofa | 100 | 100 | 100 |
| F294 | Macaca fascicularis | NA | NA | 100 |
| F295 | Macaca fascicularis | 100 | NA | 100 |
| F296 | Callosciurus notatus | NA | NA | 100 |
| F296 | Macaca fascicularis | NA | NA | 100 |
| F298 | Macaca fascicularis | 99.681 | 100 | 100 |
| F298 | Sus scrofa | NA | NA | 100 |
| F299 | Macaca fascicularis | NA | NA | 100 |
| F300 | Macaca fascicularis | NA | NA | 100 |
| F301 | Macaca fascicularis | 100 | NA | 100 |
| F302 | Macaca fascicularis | NA | NA | 100 |
| F303 | Macaca fascicularis | 99.681 | NA | 100 |
| F304 | Macaca fascicularis | NA | NA | 100 |
| F306 | Canis lupus | 100 | NA | 100 |
| F306 | Macaca fascicularis | 100 | NA | 100 |
| F307 | Macaca fascicularis | 99.681 | NA | 100 |
| F308 | Eutropis multifasciata | 98.722 | 100 | NA |
| F308 | Paradoxurus hermaphroditus | 96.711 | 97.391 | 100 |
| F308 | Macaca fascicularis | NA | NA | 100 |
| F309 | Macaca fascicularis | 99.681 | NA | 100 |
| F310 | Macaca fascicularis | NA | NA | 100 |
| F311 | Macaca fascicularis | NA | NA | 100 |
| F312 | Sus scrofa | NA | 100 | 100 |
| F312 | Macaca fascicularis | NA | NA | 100 |
| F313 | Macaca fascicularis | NA | NA | 100 |
| F314 | Sus scrofa | 100 | 100 | 100 |
| F314 | Macaca fascicularis | NA | NA | 100 |
| F318 | Maxomys sp. BOLD:AAI1690 | NA | 100 | NA |
| F318 | Galeopterus variegatus | NA | NA | 97.849 |
| F318 | Macaca fascicularis | NA | NA | 100 |
| F319 | Macaca fascicularis | NA | NA | 100 |
| F319 | Rasbora borapetensis | NA | NA | 100 |
| F319 | Sus scrofa | NA | NA | 100 |
| F320 | Macaca fascicularis | NA | NA | 100 |
| F320 | Sus scrofa | NA | NA | 100 |
| G325 | Macaca fascicularis | NA | NA | 100 |
| G328 | Macaca fascicularis | NA | NA | 100 |
| G329 | Felis catus | NA | NA | 100 |
| G329 | Macaca fascicularis | NA | NA | 100 |
| G330 | Macaca fascicularis | 100 | NA | 100 |
| G331 | Macaca fascicularis | 99.681 | NA | 100 |
| G332 | Macaca fascicularis | NA | NA | 100 |
| G333 | Macaca fascicularis | NA | NA | 100 |
| G334 | Macaca fascicularis | NA | NA | 100 |
| G335 | Macaca fascicularis | NA | NA | 100 |
| G336 | Macaca fascicularis | NA | NA | 100 |
| G337 | Macaca fascicularis | NA | NA | 100 |
| G338 | Macaca fascicularis | NA | NA | 100 |
| G338 | Sus scrofa | NA | NA | 100 |
| G339 | Macaca fascicularis | NA | NA | 100 |
| G342 | Macaca fascicularis | NA | NA | 100 |
| G344 | Macaca fascicularis | NA | NA | 100 |
| G348 | Macaca fascicularis | NA | NA | 100 |
| H350 | Macaca fascicularis | NA | NA | 100 |
| H351 | Macaca fascicularis | NA | NA | 100 |
| H352 | Macaca fascicularis | NA | NA | 100 |
| H352 | Paradoxurus hermaphroditus | NA | NA | 100 |
| H352 | Sus scrofa | NA | NA | 100 |
| H353 | Macaca fascicularis | 99.681 | NA | 100 |
| H354 | Macaca fascicularis | NA | NA | 100 |
| H355 | Macaca fascicularis | 100 | NA | 100 |
| H356 | Malayopython reticulatus | 100 | 100 | NA |
| H356 | Macaca fascicularis | NA | NA | 100 |
| H357 | Macaca fascicularis | NA | NA | 100 |
| H358 | Macaca fascicularis | NA | NA | 100 |
| H361 | Macaca fascicularis | 99.681 | NA | 100 |
| H361 | Ovis sp. | NA | NA | 100 |
| H362 | Macaca fascicularis | NA | NA | 100 |
| H365 | Macaca fascicularis | NA | NA | 100 |
| H367 | Malayopython reticulatus | NA | 100 | NA |
| H368 | Macaca fascicularis | 100 | NA | 100 |
| H369 | Macaca fascicularis | NA | NA | 100 |
| H370 | Macaca fascicularis | 100 | NA | 100 |
| H371 | Macaca fascicularis | 99.681 | NA | 100 |
| H371 | Sus scrofa | 100 | NA | 100 |
| H372 | Macaca fascicularis | NA | NA | 100 |
| H373 | Macaca fascicularis | 99.681 | NA | 100 |
| H373 | Macropus sp. | NA | NA | 100 |
| H374 | Macaca fascicularis | 100 | NA | 100 |
| H375 | Macaca fascicularis | NA | NA | 100 |
| H375 | Rattus sp. | NA | NA | 100 |
| H376 | Macaca fascicularis | NA | NA | 100 |
| I378 | Macaca fascicularis | 100 | NA | 100 |
| I380 | Sus scrofa | NA | 100 | NA |
| I381 | Macaca fascicularis | 99.681 | NA | 100 |
| I383 | Macaca fascicularis | NA | NA | 100 |
| I385 | Macaca fascicularis | 100 | NA | 100 |
| I386 | Macaca fascicularis | 100 | NA | 100 |
| I387 | Macaca fascicularis | NA | NA | 100 |
| I389 | Macaca fascicularis | NA | NA | 100 |
| I392 | Macaca fascicularis | NA | NA | 100 |
| I393 | Macaca fascicularis | NA | NA | 100 |
| I393 | Sus scrofa | NA | NA | 100 |
| I394 | Macaca fascicularis | 100 | NA | 100 |
| I394 | Paradoxurus hermaphroditus | 96.711 | NA | 100 |
| I397 | Macaca fascicularis | NA | NA | 100 |
| I398 | Macaca fascicularis | 100 | NA | 100 |
| I398 | Macaca sp. | NA | 100 | NA |
| I399 | Macaca fascicularis | 100 | NA | 100 |
| I399 | Sus scrofa | NA | NA | 100 |
| I402 | Macaca fascicularis | NA | NA | 100 |
| I403 | Macaca fascicularis | NA | NA | 100 |
| I403 | Sus scrofa | NA | NA | 100 |
| I404 | Paradoxurus hermaphroditus | 96.711 | NA | 100 |
| I405 | Macaca fascicularis | NA | NA | 100 |
| J406 | Sus scrofa | 100 | NA | NA |
| J406 | Macaca fascicularis | NA | NA | 100 |
| J407 | Macaca fascicularis | NA | NA | 100 |
| J408 | Macaca fascicularis | NA | NA | 100 |
| J409 | Macaca fascicularis | 100 | NA | 100 |
| J410 | Macaca fascicularis | 100 | NA | 100 |
| J411 | Macaca fascicularis | NA | NA | 100 |
| J411 | Rusa unicolor | NA | NA | 100 |
| J412 | Macaca fascicularis | NA | NA | 100 |
| J412 | Sus scrofa | NA | NA | 100 |
| J413 | Macaca fascicularis | 99.681 | NA | 100 |
| J414 | Macaca fascicularis | NA | NA | 100 |
| J415 | Macaca fascicularis | NA | NA | 100 |
| J416 | Macaca fascicularis | NA | NA | 100 |
| J417 | Macaca fascicularis | 100 | NA | 100 |
| J419 | Macaca fascicularis | NA | NA | 100 |
| J420 | Macaca fascicularis | 99.681 | NA | 100 |
| J421 | Rusa unicolor | NA | NA | 100 |
| J422 | Macaca fascicularis | NA | NA | 100 |
| J423 | Macaca fascicularis | NA | NA | 100 |
| J424 | Macaca fascicularis | NA | NA | 100 |
| J425 | Dama dama | NA | 100 | NA |
| J425 | Macaca fascicularis | NA | NA | 100 |
| J429 | Sus scrofa | NA | NA | 100 |
| J431 | Macaca sp. | 95.833 | NA | NA |
| J431 | Macaca fascicularis | NA | NA | 100 |
| J435 | Macaca fascicularis | 99.681 | NA | 100 |

**Supplementary Table 5: Percentage identities for vertebrate matches for the analysis of iDNA metabarcoding using Illumina (1 replicate) and MinION (1 replicate). The best percentage for a vertebrate species are reported.**

| **Sample** | **Vertebrate species** | **Illumina 1 replicate** | | | **MinION 1 replicate** | | |
| --- | --- | --- | --- | --- | --- | --- | --- |
|  |  | **244 bp COI** | **313 bp COI** | **16S** | **244bp** | **313bp** | **16S** |
| A001 | Macaca fascicularis | NA | NA | 100 | NA | NA | 100 |
| A002 | Sus scrofa | 100 | NA | NA | NA | NA | NA |
| A003 | Macaca fascicularis | NA | NA | NA | NA | NA | 100 |
| A004 | Macaca fascicularis | NA | NA | 100 | NA | NA | 99.145 |
| A005 | Macaca fascicularis | NA | NA | 100 | NA | NA | 100 |
| A006 | Sus scrofa | 100 | NA | NA | NA | NA | NA |
| A008 | Macaca fascicularis | NA | NA | 100 | NA | NA | 100 |
| A008 | Sus scrofa | NA | NA | NA | NA | NA | 100 |
| A009 | Macaca fascicularis | NA | NA | 100 | NA | NA | 100 |
| A009 | Sus scrofa | 100 | NA | NA | NA | NA | NA |
| A010 | Macaca fascicularis | NA | NA | 100 | NA | NA | 100 |
| A011 | Macaca fascicularis | NA | NA | 100 | NA | NA | 100 |
| A012 | Macaca fascicularis | NA | NA | NA | NA | NA | 100 |
| A014 | Sus scrofa | 100 | NA | NA | NA | NA | NA |
| A015 | Sus scrofa | 100 | NA | NA | 98.77 | NA | NA |
| A016 | Macaca fascicularis | NA | NA | 100 | NA | NA | 100 |
| A017 | Macaca fascicularis | NA | NA | 100 | NA | NA | NA |
| A017 | Rusa unicolor | NA | NA | 100 | NA | NA | NA |
| A017 | Sus scrofa | 100 | NA | NA | NA | NA | NA |
| A017 | Cervidae sp. | NA | NA | NA | NA | NA | 100 |
| A018 | Macaca fascicularis | NA | NA | 100 | NA | NA | 100 |
| B078 | Macaca fascicularis | NA | 100 | 100 | NA | 99.045 | 100 |
| B080 | Macaca fascicularis | NA | NA | 100 | NA | NA | 100 |
| B081 | Presbytis femoralis | NA | NA | 100 | NA | NA | NA |
| B081 | Trachypithecus francoisi | NA | NA | NA | NA | NA | 98.969 |
| B082 | Macaca fascicularis | NA | NA | 100 | NA | NA | NA |
| B082 | Macaca mulatta | NA | NA | NA | NA | NA | 100 |
| B084 | Macaca fascicularis | NA | NA | 100 | NA | NA | 100 |
| B085 | Macaca mulatta | 97.391 | 99.045 | NA | NA | NA | NA |
| B085 | Macaca fascicularis | NA | NA | 100 | NA | NA | 100 |
| B086 | Macaca fascicularis | NA | NA | 100 | NA | NA | 100 |
| B088 | Macaca fascicularis | NA | NA | 100 | NA | NA | 100 |
| B089 | Macaca fascicularis | NA | NA | 100 | NA | NA | 100 |
| B091 | Macaca fascicularis | NA | NA | 100 | NA | NA | 100 |
| B092 | Colobinae sp. | NA | NA | NA | NA | NA | 98.81 |
| B093 | Macaca fascicularis | NA | NA | 100 | NA | NA | 100 |
| B094 | Macaca fascicularis | NA | NA | NA | NA | NA | 100 |
| B095 | Macaca fascicularis | NA | NA | 100 | NA | NA | 100 |
| B095 | Manis sp. | NA | NA | 100 | NA | NA | NA |
| B096 | Macaca fascicularis | NA | NA | NA | NA | NA | 100 |
| C153 | Sus scrofa | 100 | NA | NA | NA | NA | 100 |
| C153 | Macaca fascicularis | NA | NA | NA | NA | NA | 100 |
| C155 | Macaca fascicularis | NA | NA | NA | NA | NA | 100 |
| C156 | Macaca fascicularis | NA | NA | 100 | NA | NA | 100 |
| C157 | Macaca fascicularis | NA | NA | 100 | NA | NA | 100 |
| C158 | Macaca fascicularis | 100 | NA | 100 | NA | NA | 100 |
| C159 | Macaca fascicularis | NA | NA | NA | NA | NA | 100 |
| C160 | Macaca fascicularis | NA | 100 | 100 | NA | NA | 100 |
| C161 | Macaca fascicularis | NA | 99.681 | 100 | NA | 99.035 | NA |
| C162 | Macaca fascicularis | NA | NA | 100 | NA | NA | 100 |
| C163 | Macaca fascicularis | NA | NA | 100 | NA | NA | 100 |
| C164 | Macaca fascicularis | NA | 99.681 | 100 | NA | 99.681 | 100 |
| D195 | Macaca fascicularis | NA | NA | 100 | NA | NA | 100 |
| D196 | Macaca fascicularis | NA | NA | 100 | NA | NA | 100 |
| D197 | Macaca fascicularis | NA | 99.681 | 100 | NA | 99.032 | 100 |
| D197 | Sus scrofa | 100 | NA | NA | NA | NA | NA |
| D198 | Macaca fascicularis | NA | 100 | 100 | NA | 100 | 100 |
| D198 | Pan troglodytes | NA | NA | NA | NA | NA | 97.778 |
| D198 | Sus scrofa | NA | NA | NA | NA | NA | 100 |
| D199 | Macaca fascicularis | 100 | NA | 100 | NA | NA | 100 |
| D199 | Sus scrofa | 100 | NA | NA | NA | NA | NA |
| D199 | Pan troglodytes | NA | NA | NA | NA | NA | 98.889 |
| D200 | Macaca fascicularis | NA | 100 | 100 | NA | 99.361 | 100 |
| D202 | Macaca fascicularis | NA | NA | 100 | NA | NA | 100 |
| D203 | Macaca fascicularis | NA | NA | 100 | NA | NA | 100 |
| D204 | Paradoxurus hermaphroditus | NA | NA | 100 | NA | NA | NA |
| D204 | Sus scrofa | 100 | NA | NA | NA | NA | NA |
| D204 | Macaca fascicularis | NA | NA | NA | NA | NA | 100 |
| D205 | Macaca fascicularis | NA | NA | 100 | NA | NA | 100 |
| D206 | Macaca fascicularis | NA | 99.681 | 100 | NA | 98.73 | 100 |
| D206 | Sus scrofa | 100 | NA | NA | NA | NA | NA |
| D206 | Pan troglodytes | NA | NA | NA | NA | NA | 97.5 |
| A019 | Macaca fascicularis | NA | NA | NA | NA | NA | 100 |
| A020 | Macaca fascicularis | NA | NA | 100 | NA | NA | 100 |
| A021 | Macaca fascicularis | NA | NA | NA | NA | NA | 100 |
| A022 | Macaca fascicularis | NA | NA | 100 | NA | NA | 100 |
| A024 | Macaca fascicularis | NA | NA | 100 | NA | NA | 100 |
| A025 | Macaca fascicularis | NA | NA | 100 | NA | NA | 100 |
| A025 | Gallus gallus | 100 | NA | NA | NA | NA | NA |
| A026 | Macaca fascicularis | NA | 99.681 | 100 | NA | NA | 100 |
| A027 | Macaca fascicularis | NA | NA | NA | NA | NA | 100 |
| A028 | Macaca fascicularis | NA | 99.681 | 100 | NA | 99.681 | 100 |
| A029 | Macaca fascicularis | NA | NA | 100 | NA | NA | 100 |
| A029 | Gallus gallus | 100 | NA | NA | NA | NA | NA |
| A030 | Macaca fascicularis | NA | NA | 100 | NA | NA | 100 |
| A030 | Sus scrofa | NA | NA | NA | NA | NA | 100 |
| A031 | Macaca fascicularis | NA | NA | 100 | NA | NA | 100 |
| A032 | Macaca fascicularis | NA | NA | 100 | NA | NA | 100 |
| A034 | Macaca fascicularis | NA | NA | NA | NA | NA | 100 |
| A035 | Macaca fascicularis | NA | NA | 100 | NA | NA | 100 |
| A036 | Macaca fascicularis | NA | 99.681 | 100 | NA | 99.681 | 100 |
| A037 | Macaca fascicularis | NA | NA | 100 | NA | NA | 100 |
| A038 | Macaca fascicularis | NA | NA | NA | NA | NA | 100 |
| A039 | Macaca fascicularis | NA | NA | 100 | NA | NA | 100 |
| A040 | Macaca fascicularis | NA | 100 | 100 | NA | 98.083 | 100 |
| A041 | Macaca fascicularis | NA | NA | 100 | NA | NA | 100 |
| A042 | Macaca fascicularis | NA | NA | 100 | NA | NA | 100 |
| A043 | Macaca fascicularis | NA | NA | 100 | NA | NA | 100 |
| A045 | Macaca fascicularis | NA | 99.681 | 100 | NA | 97.46 | 100 |
| A045 | Macaca mulatta | 99.13 | NA | NA | 98.76 | NA | NA |
| A046 | Macaca fascicularis | NA | NA | 100 | NA | NA | 100 |
| A047 | Macaca fascicularis | NA | NA | 100 | NA | NA | 100 |
| A047 | Sus scrofa | NA | NA | NA | NA | NA | 100 |
| B098 | Macaca fascicularis | NA | NA | 100 | NA | NA | 100 |
| B099 | Macaca fascicularis | NA | NA | 100 | NA | NA | 100 |
| B100 | Macaca fascicularis | NA | NA | 100 | NA | NA | 100 |
| B100 | Sus scrofa | NA | NA | 100 | NA | NA | 100 |
| B102 | Macaca fascicularis | NA | NA | NA | NA | NA | 100 |
| B103 | Macaca fascicularis | NA | NA | 100 | NA | NA | 100 |
| B103 | Sus scrofa | NA | NA | 100 | NA | NA | 100 |
| B104 | Tupaia glis | NA | 99.681 | NA | NA | 99.363 | 96.809 |
| B104 | Macaca fascicularis | NA | NA | 100 | NA | NA | 100 |
| B105 | Macaca fascicularis | NA | 99.681 | 100 | NA | 99.681 | 100 |
| B106 | Macaca fascicularis | NA | NA | 100 | NA | NA | 100 |
| B106 | Sus scrofa | NA | NA | NA | NA | NA | 100 |
| B108 | Macaca mulatta | NA | NA | 98.936 | NA | NA | 100 |
| B109 | Macaca fascicularis | NA | NA | NA | NA | NA | 100 |
| B111 | Macaca fascicularis | NA | NA | 100 | NA | NA | 100 |
| B112 | Macaca fascicularis | NA | NA | NA | NA | NA | 100 |
| B113 | Macaca fascicularis | NA | NA | NA | NA | NA | 100 |
| B114 | Macaca fascicularis | NA | NA | NA | NA | NA | 100 |
| B115 | Macaca fascicularis | NA | NA | 100 | NA | NA | NA |
| B116 | Macaca fascicularis | NA | NA | NA | NA | NA | 100 |
| B117 | Macaca fascicularis | NA | NA | 100 | NA | NA | 100 |
| B118 | Macaca fascicularis | NA | NA | 100 | NA | NA | 100 |
| B118 | Sus scrofa | NA | NA | 100 | NA | NA | 100 |
| B119 | Macaca fascicularis | NA | NA | 100 | NA | NA | 100 |
| B122 | Sus scrofa | 100 | 100 | 100 | 100 | 99.363 | 100 |
| B122 | Macaca fascicularis | NA | NA | 100 | NA | NA | 100 |
| C167 | Macaca fascicularis | NA | 99.681 | 100 | 100 | 99.363 | 100 |
| C167 | Macaca sp. | 100 | NA | NA | NA | NA | NA |
| C168 | Macaca fascicularis | NA | NA | NA | NA | NA | 100 |
| C169 | Macaca fascicularis | NA | 99.681 | 100 | NA | 99.681 | 100 |
| C170 | Macaca fascicularis | NA | NA | 100 | NA | NA | 100 |
| C171 | Macaca fascicularis | NA | NA | NA | NA | NA | 100 |
| C172 | Macaca fascicularis | NA | NA | NA | NA | NA | 100 |
| C173 | Macaca fascicularis | NA | 99.681 | 100 | NA | 99.038 | 100 |
| C173 | Leptocoma brasiliana | 99.083 | NA | NA | 98.354 | NA | NA |
| C174 | Macaca fascicularis | NA | NA | 100 | NA | NA | 100 |
| C174 | Gallus gallus | 100 | NA | NA | NA | NA | NA |
| C174 | Gallus sp. | NA | NA | NA | 99.602 | NA | NA |
| C174 | Anser canagicus | NA | NA | NA | 98.701 | NA | NA |
| C176 | Macaca fascicularis | NA | 100 | 100 | NA | 100 | 100 |
| C177 | Macaca fascicularis | NA | NA | 100 | NA | NA | 100 |
| C178 | Macaca fascicularis | NA | 100 | 100 | NA | 100 | 100 |
| C178 | Mixornis flavicollis | NA | 95.793 | NA | NA | NA | NA |
| C179 | Macaca fascicularis | NA | NA | NA | NA | NA | 100 |
| C180 | Macaca fascicularis | NA | NA | 100 | NA | NA | 100 |
| C181 | Macaca nemestrina | NA | 99.724 | NA | NA | 97.872 | NA |
| C181 | Macaca fascicularis | NA | NA | 100 | NA | NA | 100 |
| C181 | Ovis sp. | NA | NA | 100 | NA | NA | 100 |
| C181 | Rusa unicolor | NA | NA | 100 | NA | NA | NA |
| C181 | Cervidae sp. | NA | NA | NA | NA | NA | 100 |
| C182 | Macaca fascicularis | NA | 99.681 | 100 | NA | 99.045 | 100 |
| C183 | Macaca fascicularis | NA | 99.681 | NA | NA | NA | NA |
| C184 | Macaca fascicularis | NA | 100 | 100 | NA | 100 | 100 |
| C187 | Ovis aries | NA | 100 | NA | NA | 99.042 | NA |
| C187 | Ovis sp. | NA | NA | 100 | NA | NA | 100 |
| C187 | Macaca fascicularis | NA | NA | NA | NA | NA | 100 |
| C188 | Macaca fascicularis | NA | 99.681 | 100 | NA | NA | 100 |
| C189 | Macaca fascicularis | NA | NA | NA | NA | NA | 100 |
| C189 | Paradoxurus hermaphroditus | NA | NA | NA | NA | NA | 100 |
| C189 | Sus scrofa | NA | NA | NA | NA | NA | 100 |
| C190 | Macaca fascicularis | NA | 99.681 | 100 | NA | NA | 100 |
| C191 | Macaca fascicularis | NA | NA | 100 | NA | NA | 100 |
| C191 | Sus scrofa | NA | NA | 100 | NA | NA | 100 |
| C191 | Hemidactylus frenatus | 100 | NA | NA | NA | NA | NA |
| C192 | Macaca fascicularis | NA | 99.681 | 100 | NA | 99.045 | 100 |
| C193 | Macaca fascicularis | NA | 99.681 | 100 | NA | 99.681 | 100 |
| C193 | Sus scrofa | NA | NA | NA | NA | NA | 100 |
| C194 | Macaca fascicularis | NA | 100 | 100 | NA | NA | 100 |
| D207 | Macaca fascicularis | NA | NA | 100 | NA | NA | 100 |
| D207 | Sus scrofa | NA | NA | NA | NA | NA | 100 |
| D208 | Macaca fascicularis | NA | NA | 100 | NA | NA | 100 |
| D209 | Macaca fascicularis | NA | NA | 100 | NA | NA | 100 |
| D210 | Presbytis femoralis | NA | 100 | 100 | NA | NA | 100 |
| D212 | Macaca fascicularis | NA | NA | NA | NA | NA | 100 |
| D214 | Macaca fascicularis | NA | 100 | 100 | NA | 100 | 100 |
| D215 | Felis catus | NA | 100 | NA | NA | NA | NA |
| D215 | Macaca fascicularis | NA | NA | 100 | NA | NA | 100 |
| D215 | Cervidae sp. | NA | NA | NA | NA | NA | 100 |
| D216 | Macaca fascicularis | 100 | 99.681 | 100 | 100 | 99.681 | 100 |
| D216 | Hemidactylus frenatus | 100 | NA | NA | 100 | NA | NA |
| D217 | Gallus gallus | 100 | NA | NA | NA | NA | NA |
| D217 | Macaca fascicularis | NA | NA | NA | NA | NA | 100 |
| D217 | Sus scrofa | NA | NA | NA | NA | NA | 100 |
| D220 | Macaca fascicularis | NA | NA | 100 | NA | NA | 100 |
| D221 | Macaca fascicularis | NA | NA | 100 | NA | NA | 100 |
| D223 | Gallus gallus | 100 | NA | NA | NA | NA | NA |
| D223 | Macaca fascicularis | NA | NA | NA | NA | NA | 100 |
| D224 | Macaca fascicularis | NA | NA | 100 | NA | NA | 100 |
| D224 | Paradoxurus hermaphroditus | NA | NA | 100 | NA | NA | 100 |
| D224 | Sus scrofa | NA | NA | NA | NA | NA | 100 |
| D225 | Macaca fascicularis | NA | NA | 100 | NA | NA | 100 |
| D226 | Macaca fascicularis | NA | NA | 100 | NA | NA | 100 |
| D227 | Sus scrofa | NA | 100 | 100 | NA | NA | 100 |
| D227 | Macaca fascicularis | NA | NA | 100 | NA | NA | 100 |
| D228 | Macaca fascicularis | NA | 100 | 100 | NA | 100 | 100 |
| D228 | Gallus gallus | 100 | NA | NA | NA | NA | NA |
| D229 | Macaca fascicularis | NA | NA | 100 | NA | NA | 100 |
| D230 | Macaca fascicularis | NA | NA | 100 | NA | NA | 100 |
| D231 | Macaca fascicularis | NA | NA | NA | NA | NA | 100 |
| D232 | Sus scrofa | 100 | NA | 100 | NA | NA | 100 |
| D232 | Macaca fascicularis | NA | NA | NA | NA | NA | 100 |
| D233 | Macaca fascicularis | 100 | NA | 100 | NA | NA | 100 |
| D234 | Macaca fascicularis | NA | NA | NA | NA | NA | 100 |
| D235 | Macaca fascicularis | NA | NA | 100 | NA | NA | 100 |
| A048 | Macaca fascicularis | NA | NA | 100 | NA | NA | 100 |
| A048 | Hemidactylus frenatus | 100 | NA | NA | NA | NA | NA |
| A049 | Macaca fascicularis | NA | 100 | 100 | NA | NA | 100 |
| A050 | Macaca fascicularis | NA | NA | 100 | NA | NA | 100 |
| A051 | Macaca fascicularis | NA | NA | 100 | NA | NA | 100 |
| A052 | Macaca fascicularis | NA | 99.681 | 100 | NA | NA | 100 |
| A053 | Pan troglodytes | NA | NA | NA | 100 | NA | NA |
| A054 | Macaca fascicularis | NA | NA | 100 | NA | NA | 100 |
| A055 | Macaca fascicularis | NA | NA | 100 | NA | NA | 100 |
| A056 | Paradoxurus hermaphroditus | NA | NA | NA | NA | NA | 100 |
| A057 | Macaca fascicularis | NA | NA | 100 | NA | NA | 100 |
| A058 | Macaca fascicularis | NA | NA | 100 | NA | NA | 100 |
| A059 | Sus scrofa | NA | NA | NA | NA | NA | 100 |
| A060 | Macaca fascicularis | NA | NA | 100 | NA | NA | 100 |
| A061 | Macaca fascicularis | NA | NA | 100 | NA | NA | 100 |
| A061 | Sus scrofa | NA | NA | NA | NA | NA | 100 |
| A063 | Macaca fascicularis | NA | NA | 100 | NA | NA | 100 |
| A064 | Macaca fascicularis | NA | NA | 100 | NA | NA | 100 |
| A067 | Macaca fascicularis | NA | 99.681 | 100 | NA | 100 | 100 |
| A067 | Sus scrofa | NA | NA | NA | NA | NA | 100 |
| A068 | Macaca fascicularis | NA | 100 | 100 | NA | 99.042 | 100 |
| A068 | Rusa unicolor | 100 | NA | NA | NA | NA | NA |
| B126 | Macaca fascicularis | NA | NA | 100 | NA | NA | 100 |
| B128 | Sus scrofa | 100 | NA | NA | NA | NA | NA |
| B129 | Macaca fascicularis | NA | NA | 100 | NA | NA | 100 |
| B129 | Sus scrofa | 100 | NA | NA | NA | NA | 100 |
| B130 | Galeopterus variegatus | NA | NA | 97.849 | NA | NA | 98.925 |
| B130 | Macaca fascicularis | NA | NA | 100 | NA | NA | 100 |
| B132 | Macaca fascicularis | NA | NA | NA | NA | NA | 100 |
| B133 | Sus scrofa | 100 | NA | NA | NA | NA | NA |
| B133 | Macaca fascicularis | NA | NA | NA | NA | NA | 100 |
| B134 | Sus scrofa | 100 | NA | 100 | 100 | NA | 100 |
| B134 | Macaca fascicularis | NA | NA | NA | NA | NA | 100 |
| B135 | Macaca fascicularis | 100 | 100 | 100 | NA | 100 | 100 |
| B135 | Pycnonotus plumosus | NA | 99.042 | NA | NA | 98.701 | NA |
| B135 | Suncus sp. | NA | NA | 100 | NA | NA | 100 |
| B135 | Pan troglodytes | NA | NA | NA | NA | NA | 97.778 |
| B135 | Sus scrofa | NA | NA | NA | NA | NA | 100 |
| B136 | Macaca fascicularis | NA | NA | 100 | NA | NA | 100 |
| B137 | Sus scrofa | NA | NA | 100 | NA | NA | 100 |
| B141 | Macaca mulatta | NA | NA | 100 | NA | NA | 100 |
| B141 | Sus scrofa | 100 | NA | NA | NA | NA | NA |
| B143 | Sus scrofa | 100 | NA | NA | NA | NA | NA |
| B144 | Macaca fascicularis | NA | NA | NA | NA | NA | 100 |
| B145 | Macaca fascicularis | NA | NA | 100 | NA | NA | 100 |
| B146 | Macaca fascicularis | NA | NA | 100 | NA | NA | 100 |
| E237 | Macaca fascicularis | NA | NA | 100 | NA | NA | 100 |
| E239 | Macaca fascicularis | NA | NA | 100 | NA | NA | 100 |
| E240 | Macaca fascicularis | NA | NA | 100 | NA | NA | 100 |
| E242 | Macaca fascicularis | NA | NA | 100 | NA | NA | 100 |
| E243 | Macaca fascicularis | NA | 99.681 | 100 | NA | 99.361 | 100 |
| E244 | Macaca fascicularis | NA | NA | 100 | NA | NA | 100 |
| E246 | Macaca fascicularis | NA | NA | 100 | NA | NA | 100 |
| E248 | Macaca fascicularis | NA | NA | 100 | NA | NA | 100 |
| E250 | Sus scrofa | NA | NA | 100 | NA | NA | 100 |
| E250 | Macaca fascicularis | NA | NA | NA | NA | NA | 100 |
| E251 | Macaca fascicularis | 100 | 99.681 | 100 | NA | 99.682 | 100 |
| E252 | Macaca fascicularis | NA | NA | NA | NA | NA | 100 |
| E253 | Macaca fascicularis | NA | NA | 100 | NA | NA | 100 |
| E254 | Macaca fascicularis | NA | NA | 100 | NA | NA | 100 |
| E257 | Macaca mulatta | NA | NA | 100 | NA | NA | 100 |
| E257 | Macaca sp. | 100 | NA | NA | NA | NA | NA |
| E257 | Macaca fascicularis | NA | NA | NA | 100 | NA | NA |
| E257 | Pan troglodytes | NA | NA | NA | NA | NA | 98.901 |
| E257 | Sus scrofa | NA | NA | NA | NA | NA | 100 |
| E258 | Gallus gallus | 100 | NA | NA | NA | NA | NA |
| E258 | Pan troglodytes | NA | NA | NA | 99.052 | NA | NA |
| F270 | Macaca fascicularis | NA | NA | 100 | NA | NA | 100 |
| F271 | Macaca fascicularis | NA | NA | 100 | NA | NA | 100 |
| F272 | Macaca fascicularis | NA | NA | 100 | NA | NA | NA |
| F273 | Macaca fascicularis | NA | NA | 100 | NA | NA | 100 |
| F274 | Macaca fascicularis | NA | NA | NA | NA | NA | 100 |
| F275 | Macaca fascicularis | NA | NA | 100 | NA | NA | 100 |
| F276 | Macaca fascicularis | NA | NA | 100 | NA | NA | 100 |
| F277 | Macaca fascicularis | NA | NA | 100 | NA | NA | 100 |
| F279 | Sus scrofa | 100 | NA | NA | NA | NA | NA |
| F280 | Macaca fascicularis | NA | NA | 100 | NA | NA | 100 |
| F280 | Rusa unicolor | 100 | NA | 100 | 100 | NA | NA |
| F280 | Sus scrofa | 100 | NA | 100 | 99.59 | NA | 100 |
| F280 | Cervidae sp. | NA | NA | NA | NA | NA | 100 |
| F281 | Macaca fascicularis | NA | NA | NA | NA | NA | 100 |
| F282 | Macaca fascicularis | NA | NA | 100 | NA | NA | 100 |
| F283 | Macaca fascicularis | NA | NA | 100 | NA | NA | 100 |
| F284 | Macaca fascicularis | NA | NA | NA | NA | NA | 100 |
| F285 | Macaca fascicularis | NA | NA | 100 | NA | NA | 100 |
| F286 | Macaca fascicularis | NA | NA | NA | NA | NA | 99.138 |
| F289 | Macaca fascicularis | NA | 100 | 100 | NA | NA | 100 |
| F290 | Macaca fascicularis | NA | NA | 100 | NA | NA | 100 |
| A069 | Macaca fascicularis | NA | NA | NA | NA | NA | 100 |
| A072 | Sus scrofa | NA | NA | NA | NA | NA | 100 |
| A073 | Macaca fascicularis | NA | NA | 100 | NA | NA | 100 |
| A074 | Macaca fascicularis | NA | NA | 100 | NA | NA | 100 |
| A075 | Macaca fascicularis | NA | NA | 100 | NA | NA | 100 |
| A075 | Sus scrofa | NA | NA | NA | NA | NA | 100 |
| A077 | Macaca fascicularis | NA | 100 | 100 | NA | 100 | 100 |
| B147 | Macaca fascicularis | NA | 100 | 100 | NA | 99.681 | 100 |
| B148 | Macaca fascicularis | NA | NA | 100 | NA | NA | 100 |
| B149 | Macaca fascicularis | NA | NA | 100 | NA | NA | 100 |
| B149 | Sus scrofa | NA | NA | 100 | NA | NA | 100 |
| B150 | Macaca fascicularis | NA | NA | NA | NA | NA | 100 |
| B151 | Macaca fascicularis | NA | NA | 100 | NA | NA | 100 |
| B152 | Megalobrama sp. | NA | 100 | NA | NA | 98.722 | NA |
| E259 | Macaca fascicularis | NA | 100 | 100 | NA | NA | 100 |
| E259 | Sus scrofa | NA | NA | NA | NA | NA | 100 |
| E261 | Macaca fascicularis | NA | 99.681 | 100 | NA | NA | 100 |
| E261 | Sus scrofa | 100 | 99.361 | 100 | 100 | NA | 100 |
| E263 | Macaca fascicularis | NA | NA | 100 | NA | NA | 100 |
| E263 | Macaca mulatta | 98.261 | NA | NA | NA | NA | NA |
| E265 | Sus scrofa | 100 | NA | 100 | NA | NA | 100 |
| E265 | Macaca fascicularis | NA | NA | NA | NA | NA | 100 |
| E266 | Macaca fascicularis | 100 | 100 | 100 | NA | 100 | 100 |
| E266 | Sus scrofa | NA | NA | NA | NA | NA | 100 |
| E267 | Callosciurus notatus | NA | NA | 100 | NA | NA | 100 |
| E267 | Sus scrofa | 100 | NA | 100 | 100 | NA | 100 |
| E267 | Macaca fascicularis | NA | NA | NA | NA | NA | 100 |
| E268 | Macaca fascicularis | NA | NA | 100 | NA | NA | 100 |
| F291 | Macaca fascicularis | NA | NA | 100 | NA | NA | 100 |
| F292 | Presbytis femoralis | NA | 100 | 100 | NA | 99.042 | 100 |
| F292 | Sus scrofa | 100 | 100 | 100 | 100 | 100 | 100 |
| F294 | Macaca fascicularis | NA | NA | 100 | NA | NA | 100 |
| F295 | Macaca fascicularis | NA | 100 | 100 | NA | 100 | 100 |
| F296 | Macaca fascicularis | NA | 99.681 | 100 | NA | 97.771 | 100 |
| F296 | Callosciurus notatus | NA | NA | 100 | NA | NA | NA |
| F298 | Macaca fascicularis | 100 | 99.681 | 100 | 100 | 99.363 | 100 |
| F298 | Sus scrofa | NA | NA | NA | NA | NA | 100 |
| F299 | Macaca fascicularis | NA | NA | 100 | NA | NA | 100 |
| F300 | Macaca fascicularis | NA | NA | 100 | NA | NA | 100 |
| F301 | Macaca fascicularis | NA | 100 | 100 | NA | NA | NA |
| F301 | Macaca mulatta | NA | NA | NA | NA | 98.413 | 100 |
| F301 | Sus scrofa | NA | NA | NA | NA | NA | 100 |
| F302 | Macaca fascicularis | NA | NA | 100 | NA | NA | 100 |
| F303 | Macaca fascicularis | NA | 99.681 | 100 | NA | 99.681 | 100 |
| F303 | Sus scrofa | NA | NA | NA | NA | NA | 100 |
| F304 | Macaca fascicularis | NA | NA | 100 | NA | NA | 100 |
| F306 | Canis lupus | NA | 100 | 100 | NA | NA | 100 |
| F306 | Macaca fascicularis | NA | 100 | 100 | NA | 99.682 | 100 |
| F306 | Sus scrofa | NA | NA | NA | NA | NA | 100 |
| F307 | Macaca fascicularis | NA | 99.681 | 100 | NA | 99.681 | 100 |
| F308 | Paradoxurus hermaphroditus | 97.391 | 96.711 | 100 | 96.327 | NA | 100 |
| F308 | Eutropis multifasciata | 100 | NA | NA | NA | NA | NA |
| F308 | Macaca fascicularis | NA | NA | NA | NA | NA | 100 |
| F308 | Sus scrofa | NA | NA | NA | NA | NA | 100 |
| F309 | Macaca fascicularis | NA | 99.681 | 100 | NA | 99.681 | 100 |
| F310 | Macaca fascicularis | NA | NA | 100 | NA | NA | 100 |
| F311 | Macaca fascicularis | NA | 100 | 100 | NA | 98.726 | 100 |
| F311 | Pan troglodytes | NA | NA | NA | NA | NA | 97.701 |
| F312 | Sus scrofa | 100 | 100 | 100 | 100 | 99.363 | 100 |
| F312 | Macaca fascicularis | NA | NA | 100 | NA | NA | 100 |
| F313 | Macaca fascicularis | NA | NA | NA | NA | NA | 100 |
| F314 | Sus scrofa | NA | 100 | 100 | NA | NA | 100 |
| F314 | Macaca fascicularis | NA | NA | 100 | NA | NA | 100 |
| F317 | Macaca mulatta | NA | 99.681 | NA | NA | 97.444 | NA |
| F317 | Macaca fascicularis | NA | NA | 100 | NA | NA | 100 |
| F318 | Galeopterus variegatus | NA | NA | 97.849 | NA | NA | 98.889 |
| F318 | Macaca fascicularis | NA | NA | 100 | NA | NA | 100 |
| F318 | Maxomys sp. BOLD:AAI1690 | 100 | NA | NA | NA | NA | NA |
| F318 | Sus scrofa | NA | NA | NA | NA | NA | 100 |
| F319 | Macaca fascicularis | NA | NA | 100 | NA | NA | 100 |
| F319 | Rasbora borapetensis | NA | NA | 100 | NA | NA | NA |
| F320 | Sus scrofa | NA | 100 | 100 | NA | NA | 100 |
| F320 | Macaca fascicularis | NA | NA | 100 | NA | NA | 100 |
| G321 | Macaca fascicularis | NA | NA | NA | NA | NA | 100 |
| G323 | Macaca fascicularis | NA | NA | NA | NA | NA | 100 |
| G324 | Macaca fascicularis | NA | NA | NA | NA | NA | 100 |
| G325 | Macaca fascicularis | NA | 100 | 100 | NA | 98.046 | 100 |
| G328 | Macaca fascicularis | NA | NA | 100 | NA | NA | 100 |
| G328 | Pan troglodytes | NA | NA | NA | 100 | NA | NA |
| G329 | Felis catus | NA | NA | 100 | NA | NA | 100 |
| G329 | Macaca fascicularis | NA | NA | 100 | NA | NA | 100 |
| G330 | Macaca fascicularis | NA | 100 | 100 | NA | 99.681 | 100 |
| G331 | Macaca fascicularis | NA | 99.681 | 100 | NA | 97.134 | 100 |
| G332 | Macaca fascicularis | NA | NA | 100 | NA | NA | 100 |
| G333 | Macaca fascicularis | NA | NA | 100 | NA | NA | 100 |
| G334 | Macaca fascicularis | NA | NA | 100 | NA | NA | 100 |
| G335 | Macaca fascicularis | NA | NA | NA | NA | NA | 100 |
| G336 | Macaca fascicularis | NA | NA | 100 | NA | NA | NA |
| G337 | Macaca fascicularis | NA | NA | 100 | NA | NA | 100 |
| G338 | Macaca fascicularis | NA | NA | 100 | NA | NA | 100 |
| G338 | Sus scrofa | NA | NA | 100 | NA | NA | NA |
| G339 | Macaca fascicularis | NA | NA | 100 | NA | NA | 100 |
| G340 | Macaca fascicularis | NA | NA | NA | NA | NA | 100 |
| G340 | Tupaia glis | NA | NA | NA | NA | NA | 96.739 |
| G342 | Macaca fascicularis | NA | NA | 100 | NA | NA | 100 |
| G344 | Macaca fascicularis | NA | NA | 100 | NA | NA | 100 |
| G344 | Pan troglodytes | NA | NA | NA | 99.531 | NA | NA |
| G348 | Macaca fascicularis | NA | NA | 100 | NA | NA | 100 |
| H350 | Macaca fascicularis | NA | NA | 100 | NA | NA | 100 |
| H351 | Macaca fascicularis | NA | NA | 100 | NA | NA | 100 |
| H352 | Sus scrofa | NA | NA | 100 | NA | NA | 100 |
| H352 | Macaca fascicularis | NA | NA | NA | NA | NA | 100 |
| H352 | Paradoxurus hermaphroditus | NA | NA | NA | NA | NA | 100 |
| H353 | Macaca fascicularis | NA | 99.681 | 100 | NA | 98.726 | 100 |
| H354 | Macaca fascicularis | NA | NA | 100 | NA | NA | 100 |
| H355 | Macaca fascicularis | NA | 100 | 100 | NA | NA | 100 |
| H356 | Malayopython reticulatus | 100 | 100 | NA | 100 | NA | NA |
| H356 | Macaca fascicularis | NA | NA | 100 | NA | NA | 100 |
| H356 | Sus scrofa | NA | NA | NA | NA | NA | 100 |
| H357 | Macaca fascicularis | NA | NA | 100 | NA | NA | 100 |
| H358 | Macaca fascicularis | NA | NA | 100 | NA | NA | 100 |
| H361 | Macaca fascicularis | NA | 99.681 | 100 | NA | 99.048 | 100 |
| H361 | Ovis sp. | NA | NA | 100 | NA | NA | 100 |
| H362 | Macaca fascicularis | NA | NA | 100 | NA | NA | 100 |
| H365 | Macaca fascicularis | NA | NA | 100 | NA | NA | 100 |
| H366 | Gallus gallus | 100 | NA | NA | NA | NA | NA |
| H367 | Malayopython reticulatus | 100 | NA | NA | NA | NA | NA |
| H368 | Macaca fascicularis | NA | NA | 100 | NA | NA | 100 |
| H369 | Macaca fascicularis | NA | NA | 100 | NA | NA | 100 |
| H370 | Felis silvestris,Felis catus | NA | 99.042 | NA | NA | NA | NA |
| H370 | Macaca fascicularis | NA | 100 | 100 | NA | NA | 100 |
| H371 | Macaca fascicularis | NA | 100 | 100 | NA | 99.048 | 100 |
| H371 | Sus scrofa | NA | 100 | 100 | NA | NA | 100 |
| H372 | Macaca fascicularis | NA | NA | 100 | NA | NA | 100 |
| H373 | Macaca fascicularis | NA | 99.681 | 100 | NA | 99.679 | 100 |
| H374 | Macaca fascicularis | NA | 100 | 100 | NA | 99.351 | 100 |
| H375 | Macaca fascicularis | NA | NA | 100 | NA | NA | 100 |
| H376 | Macaca fascicularis | NA | NA | 100 | NA | NA | 100 |
| I378 | Macaca fascicularis | NA | NA | 100 | NA | NA | 100 |
| I379 | Macaca fascicularis | NA | NA | NA | NA | NA | 100 |
| I380 | Sus scrofa | 100 | NA | 98.936 | 96.735 | NA | 100 |
| I380 | Macaca fascicularis | NA | NA | NA | NA | NA | 100 |
| I381 | Macaca fascicularis | 100 | 99.683 | 100 | 100 | 99.681 | 100 |
| I383 | Macaca fascicularis | NA | NA | NA | NA | NA | 100 |
| I385 | Macaca fascicularis | NA | 100 | 100 | NA | 99.042 | 100 |
| I386 | Macaca fascicularis | NA | 100 | 100 | NA | NA | 100 |
| I386 | Pan troglodytes | NA | NA | NA | NA | NA | 97.561 |
| I387 | Macaca fascicularis | NA | NA | NA | NA | NA | 100 |
| I389 | Macaca fascicularis | NA | NA | 100 | NA | NA | 100 |
| I392 | Macaca fascicularis | NA | NA | 100 | NA | NA | 100 |
| I393 | Macaca fascicularis | NA | NA | 100 | NA | NA | 100 |
| I393 | Sus scrofa | NA | NA | 100 | NA | NA | NA |
| I394 | Macaca fascicularis | NA | 100 | 100 | NA | NA | 100 |
| I394 | Paradoxurus hermaphroditus | NA | NA | 100 | NA | NA | 100 |
| I395 | Sus scrofa | NA | NA | NA | NA | NA | 100 |
| I397 | Macaca fascicularis | NA | NA | 100 | NA | NA | 100 |
| I397 | Sus scrofa | NA | NA | 100 | NA | NA | 100 |
| I398 | Macaca fascicularis | NA | 100 | 100 | NA | 99.361 | 100 |
| I398 | Macaca sp. | 100 | NA | NA | NA | NA | NA |
| I399 | Macaca fascicularis | NA | 100 | 100 | NA | NA | 100 |
| I399 | Sus scrofa | NA | NA | 100 | NA | NA | 100 |
| I401 | Macaca fascicularis | NA | NA | NA | NA | NA | 100 |
| I402 | Macaca fascicularis | NA | NA | 100 | NA | NA | NA |
| I402 | Macaca mulatta | NA | NA | NA | NA | NA | 98.98 |
| I403 | Macaca fascicularis | NA | NA | 100 | NA | NA | 100 |
| I404 | Paradoxurus hermaphroditus | NA | 96.711 | 100 | NA | 96.711 | NA |
| I404 | Sus scrofa | NA | NA | 100 | NA | NA | 100 |
| I405 | Macaca fascicularis | NA | NA | NA | NA | NA | 100 |
| J406 | Sus scrofa | NA | 100 | NA | NA | NA | NA |
| J406 | Macaca fascicularis | NA | NA | 100 | NA | NA | 100 |
| J407 | Macaca fascicularis | NA | NA | 100 | NA | NA | 100 |
| J408 | Macaca fascicularis | NA | NA | 100 | NA | NA | 100 |
| J409 | Macaca fascicularis | NA | 100 | 100 | NA | 98.083 | 100 |
| J409 | Sundamys annandalei | 100 | NA | NA | NA | NA | NA |
| J409 | Pan troglodytes | NA | NA | NA | NA | NA | 98.889 |
| J410 | Macaca fascicularis | NA | 100 | 100 | NA | 99.361 | 100 |
| J411 | Macaca fascicularis | NA | NA | 100 | NA | NA | 100 |
| J411 | Rusa unicolor | NA | NA | 100 | NA | NA | NA |
| J411 | Cervidae sp. | NA | NA | NA | NA | NA | 100 |
| J412 | Sus scrofa | 100 | NA | 100 | 98.776 | NA | 100 |
| J412 | Macaca fascicularis | NA | NA | NA | NA | NA | 100 |
| J413 | Macaca fascicularis | NA | 99.681 | 100 | NA | 99.681 | 100 |
| J414 | Macaca fascicularis | NA | 99.681 | 100 | NA | 99.045 | 100 |
| J414 | Rusa unicolor | NA | NA | 100 | NA | NA | NA |
| J414 | Cervidae sp. | NA | NA | NA | NA | NA | 100 |
| J415 | Macaca fascicularis | NA | NA | 100 | NA | NA | 100 |
| J416 | Macaca fascicularis | NA | NA | NA | NA | NA | 100 |
| J417 | Macaca fascicularis | NA | 100 | 100 | NA | 100 | 100 |
| J419 | Macaca fascicularis | NA | NA | 100 | NA | NA | 100 |
| J420 | Macaca fascicularis | NA | 99.681 | 100 | NA | 99.361 | 100 |
| J421 | Macaca fascicularis | NA | NA | 100 | NA | NA | 100 |
| J421 | Rusa unicolor | NA | NA | 100 | NA | NA | NA |
| J421 | Cervidae sp. | NA | NA | NA | NA | NA | 100 |
| J422 | Macaca fascicularis | NA | NA | 100 | NA | NA | 100 |
| J423 | Macaca fascicularis | NA | NA | 100 | NA | NA | 100 |
| J425 | Macaca fascicularis | NA | NA | 100 | NA | NA | 100 |
| J425 | Dama dama | 100 | NA | NA | NA | NA | NA |
| J428 | Macaca fascicularis | NA | NA | 100 | NA | NA | 100 |
| J428 | Cervidae sp. | NA | NA | NA | NA | NA | 100 |
| J428 | Sus scrofa | NA | NA | NA | NA | NA | 100 |
| J429 | Sus scrofa | NA | NA | 100 | NA | NA | 100 |
| J430 | Macaca fascicularis | NA | NA | NA | NA | NA | 100 |
| J431 | Macaca nemestrina | NA | 95.86 | NA | NA | 95.847 | NA |
| J431 | Macaca fascicularis | NA | NA | 100 | NA | NA | 100 |
| J432 | Macaca fascicularis | NA | NA | NA | NA | NA | 100 |
| J433 | Macaca fascicularis | NA | NA | NA | NA | NA | 100 |
| J435 | Macaca fascicularis | NA | 99.681 | 100 | NA | NA | 100 |

**Supplementary Material 1:** Logistic regression model to determine whether type of sample, sex and family of fly can explain the probability of presence of verte

```
Logistic Model with Interactions

Call:
glm(formula = Vertpresence ~ Type.of.sample * Sex * Family, family = binomial(link = "logit"),
    data = r)

Deviance Residuals:
    Min       1Q   Median       3Q      Max
-1.8158  -1.1545   0.6536   1.0142   1.7552

Coefficients: (5 not defined because of singularities)
                                               Estimate Std. Error z value Pr(>|z|)
(Intercept)                                     1.43508    0.24881   5.768 8.03e-09 ***
Type.of.sampleBrown                            -0.18232    0.52554  -0.347  0.72865
Type.of.sampleClear                            -1.03820    0.32371  -3.207  0.00134 **
Type.of.sampleEmpty                            -1.72277    0.80327  -2.145  0.03198 *
SexM                                           -0.58779    0.46972  -1.251  0.21080
FamilyMuscidae                                 -2.73437    0.69724  -3.922 8.79e-05 ***
FamilySarcophagidae                            -0.74194    1.24976  -0.594  0.55274
Type.of.sampleBrown:SexM                        0.43364    1.32976   0.326  0.74435
Type.of.sampleClear:SexM                        0.13684    0.60968   0.224  0.82241
Type.of.sampleEmpty:SexM                      -14.69060 1029.12186  -0.014  0.98861
Type.of.sampleBrown:FamilyMuscidae              1.73292    0.97693   1.774  0.07609 .
Type.of.sampleClear:FamilyMuscidae              2.04980    0.85057   2.410  0.01596 *
Type.of.sampleEmpty:FamilyMuscidae              2.66537    1.14557   2.327  0.01998 *
Type.of.sampleBrown:FamilySarcophagidae       -31.89428 1782.49262  -0.018  0.98572
Type.of.sampleClear:FamilySarcophagidae         0.34506    1.89863   0.182  0.85579
Type.of.sampleEmpty:FamilySarcophagidae         0.33647    1.90925   0.176  0.86011
SexM:FamilyMuscidae                             0.04548    1.35850   0.033  0.97329
SexM:FamilySarcophagidae                       15.97153 1029.12345   0.016  0.98762
Type.of.sampleBrown:SexM:FamilyMuscidae              NA         NA      NA       NA
Type.of.sampleClear:SexM:FamilyMuscidae              NA         NA      NA       NA
Type.of.sampleEmpty:SexM:FamilyMuscidae              NA         NA      NA       NA
Type.of.sampleBrown:SexM:FamilySarcophagidae         NA         NA      NA       NA
Type.of.sampleClear:SexM:FamilySarcophagidae    0.04548 1455.39967   0.000  0.99998
Type.of.sampleEmpty:SexM:FamilySarcophagidae         NA         NA      NA       NA
---
Signif. codes:  0 ‘***’ 0.001 ‘**’ 0.01 ‘*’ 0.05 ‘.’ 0.1 ‘ ’ 1

(Dispersion parameter for binomial family taken to be 1)

    Null deviance: 519.68  on 391  degrees of freedom
Residual deviance: 468.33  on 373  degrees of freedom
AIC: 506.33

Number of Fisher Scoring iterations: 14
```

**Supplementary Figure 1:** Number of flies giving vertebrate identifications classified by type of sample. Individual bars represent males and females and each group represents the number of vertebrate identifications in each fly.

|  | Brown | Clear | Brown+clear | Empty |
| --- | --- | --- | --- | --- |
| 0 F | 13 | 51 | 32 | 16 |
| 0 M | 2 | 21 | 9 | 3 |
| 1 F | 24 | 57 | 72 | 10 |
| 1 M | 2 | 19 | 15 | 1 |
| 2 F | 4 | 12 | 15 | 1 |
| 2 M | 1 | 2 | 6 | 0 |
| 3 F | 2 | 0 | 2 | 0 |
| 3 M | 0 | 0 | 0 | 0 |

**Supplementary Figure 2:** Distribution of H_2_’ (Shannon entropy) values for null models. Observed value is shown by the red line.

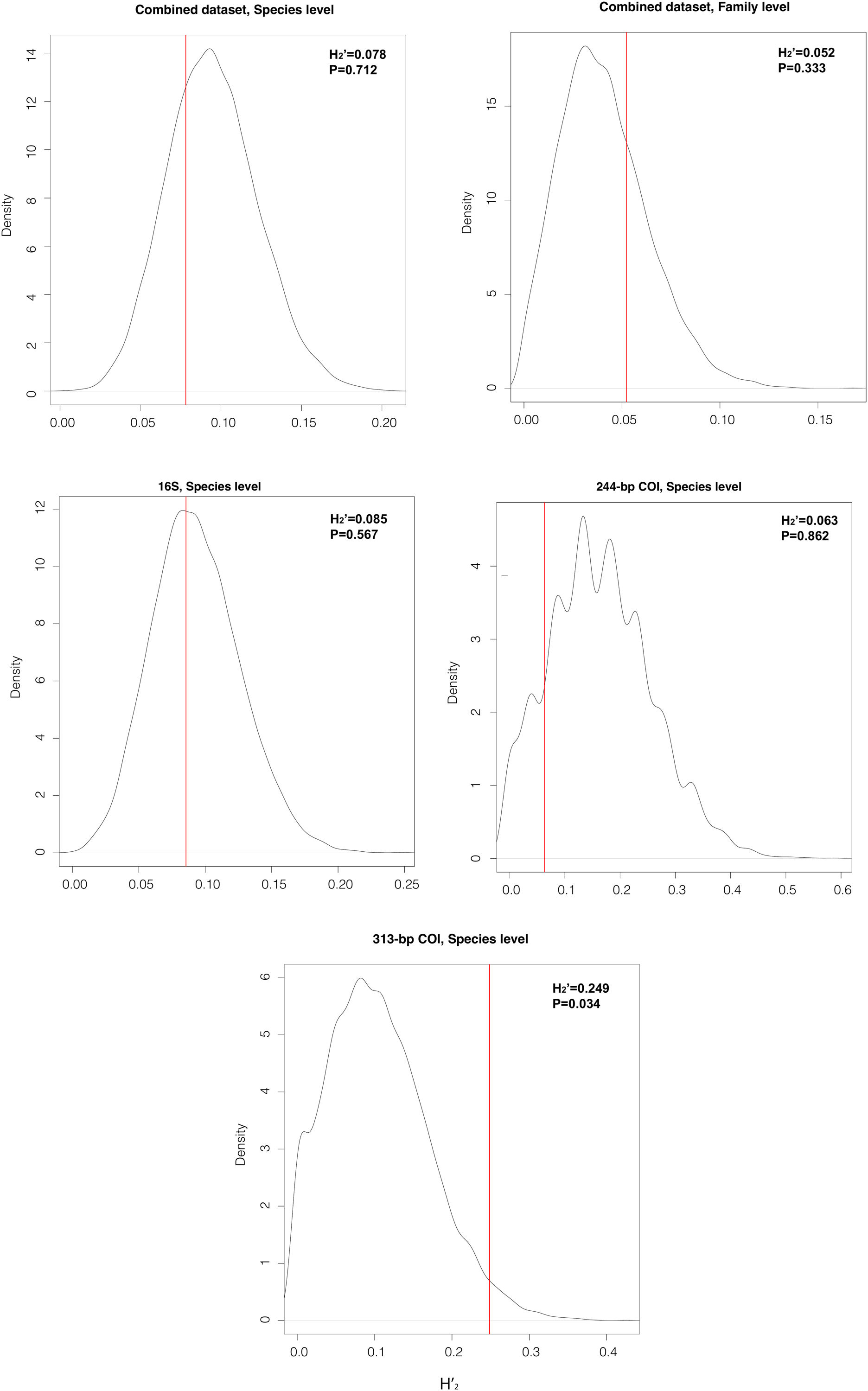
